# Benchmarking the Impact of Data Leakage on the Performance of Knowledge Graph Embedding Models for Biomedical Link Prediction

**DOI:** 10.1101/2025.01.23.634511

**Authors:** Galadriel Brière, Thomas Stosskopf, Benjamin Loire, Anaïs Baudot

## Abstract

**Motivation:** Knowledge Graphs (KGs) organize complex biomedical knowledge into structured representations of entities and relations. Knowledge Graph Embedding (KGE) models learn compact representations of KGs, and are widely applied for biomedical link prediction. Despite extensive work on KGE models, current evaluations often overlook the issue of data leakage, which can artificially inflate performance and undermine benchmark validity. Data leakage can arise when (1) there is redundancy between training and test sets, (2) the model leverages illegitimate features, or (3) the test set does not accurately reflect real-world inference.

**Results:** We assess the impact of data leakage on KGE-based link prediction across three biomedical knowledge graphs, using decoder-only and GNN-based models. We first demonstrate the impact of train-test redundancies and implement a systematic procedure to detect and remove them. Using permutation experiments, we also investigate whether node degree acts as an illegitimate predictive feature, and find no evidence that predictions are driven by degree alone. Finally, we evaluate how well common test set sampling strategies reflect real-world inference in drug repurposing. We compare random and cold-start data splits with an independent test set from Orphanet, and observe a substantial performance drop on the latter, indicating that current benchmarking practices may overestimate how well KGE models generalize to practical applications. Overall, our findings highlight the importance of rigorous benchmark design and careful evaluation of the generalization ability of KGE models for biomedical link prediction.

**Availability and Implementation:** Code and results are openly available on GitHub at https://github.com/galadrielbriere/data_leakage_kge_benchmark.git.

**Supplementary Information:** Supplementary data are available.

## Introduction

Biomedical knowledge spans multiple scales of biological organization, from molecular interactions (gene regulatory networks, protein–protein interactions, metabolic pathways) to cellular coordination, tissue and organ-level processes, and clinical relationships between diseases, phenotypes, and therapeutics. Domain-specific ontologies such as the Gene Ontology (1) and Human Phenotype Ontology (2) provide standardized terminologies for this complex knowledge. Taken altogether, this large-scale and heterogeneous domain knowledge coalesce into Knowledge Graphs (KGs), which represent entities and their relationships using triples in the form (*head, relation, tail*). KGs provide an intuitive and unified representation of multi-scale biological knowledge. By combining the structured knowledge stored in KGs with Machine Learning (ML) approaches, Knowledge Graph Embedding (KGE) methods encode entities and relations into continuous, low-dimensional latent vector spaces while preserving the fundamental structural, topological, and semantic characteristics of KGs. These latent representations facilitate data exploration and enable efficient inference of novel interactions (3), a task known as link prediction.

Drug repurposing, which involves finding new therapeutic uses for existing drugs, stands out as a critical application of link prediction where KGE can have a significant impact (4). In a knowledge graph perspective, this translates to predicting missing indication triples (triples of the form (drug, indication, disease)) that connect drugs to the diseases they can treat. By leveraging the ability of KGE to predict such novel indications, researchers can identify potential drug candidates more efficiently, reducing the time and cost associated with traditional drug discovery processes.

Despite the significant potential of KGE-based link prediction in biomedicine, challenges remain in evaluating the reliability and generalizability of these models. One key issue is data leakage. Data leakage can artificially inflate any ML predictor performance and undermine the validity of the evaluation process. Data leakage is defined as “*the relationship between (i) the reported performance scores of an ML-based predictor, (ii) the data used to train or validate the predictor, (iii) the data used to generate the reported performances of the ML-based predictor and (iv) the data on which the ML-based predictor should be run at inference time once it is deployed in its intended use case*” (5; 6).

Data leakage can be caused by multiple mechanisms. First, improper split of KG triples can result in data redundancy between the training and test sets, compromising the separation required between these datasets. *Akrami et al*. (7) demonstrated this issue through the study of widely-used nonbiomedical benchmark KGs (*FB15k, WN18*, and *YAGO3-10* ). Their findings revealed that eliminating these redundancies caused a significant decrease in the reported performance of the KGE models. The authors concluded that, under realistic conditions, KGE models do not provide an effective solution for the task of link prediction.

Second, ML models may exploit illegitimate features as prediction shortcuts. In sequence-based PPI prediction, Bernett et al. (8) showed that deep learning models relied on unintended topological features like node degree rather than protein sequences. By randomizing training networks via degree-preserving permutations, they demonstrated that models retained predictive power despite removing biologically meaningful edges, revealing reliance on structural shortcuts over biological features. While not yet demonstrated for KGE, graph structure encoded in triples could similarly enable models to exploit properties like node degree for link prediction. Investigating whether KGE models rely on such structural shortcuts is crucial to avoid overestimating their true predictive capabilities.

Finally, data leakage can arise when the training and test sets share similar data distributions that differ from the distribution encountered at deployment. This form of leakage may stem from sampling bias: test sets that fail to reflect real-world inference conditions yield inflated and unreliable performance metrics (5). In biomedical KGs, entities vary widely in connectivity, and randomly sampled test triples inevitably mirror the connectivity distribution of the training data. Yet real-world inference often concerns sparsely connected entities, such as rare diseases or underrepresented compounds, whose connectivity patterns may differ markedly from those seen during training. This discrepancy underscores the need for evaluation protocols that more accurately model realistic inference scenarios by designing test sets that better match deployment conditions and limit unintended similarities with the training data.

To the best of our knowledge, existing benchmarks for biomedical KG analysis have not examined data leakage issues thoroughly. Some benchmarks implemented measures to mitigate contamination of the test set by redundancies with the training set. For instance, (9) ensured that known reverse relations, such as “is a” and “is a reverse,” were allocated within the same split. In another example, (10) proposed an evaluation protocol based on “cold-start” triples, which ensured that the models were evaluated on entirely new and unseen combinations of entities and relations. Similarly, (11) presented an evaluation framework termed “free-setting”, in which all triples involving the relation to be predicted were removed from the training set. Lastly, (12) highlighted potential biases in inferring links for underrepresented entities in biomedical KGs, but these biases were not empirically assessed for their impact on model performance. Overall, while these efforts addressed certain aspects of data leakage, they did not systematically identify and mitigate the full range of leakage sources. Addressing this gap is essential for improving the robustness and reliability of models and benchmarks in the biomedical domain.

In this study, we propose strategies to systematically evaluate the impact of the three sources of data leakage described in (5). Our framework intends to enable comprehensive assessments of KGE models’ performance under controlled data leakage conditions, with particular emphasis on drug repurposing scenarios. We develop specific experimental protocols for evaluating each source of data leakage:

- **Data Leakage 1 (DL1): Redundancy between Training and Test Sets**. We introduce a mitigation procedure to remove redundancies between splits. Comparing KGE performance with and without this procedure quantifies artificial inflation due to redundant information.
- **Data Leakage 2 (DL2): Use of Illegitimate Features**. We investigate whether KGE models exploit node degree as a prediction proxy by training on both the original KG and a degree-preserving permuted version where indication triples are randomly shuffled Performance comparison reveals the extent to which degree alone drives predictions, despite the disruption of meaningful biological associations.
- **Data Leakage 3 (DL3): Sampling Bias in Test Set Construction**. We examine how test set sampling strategies can lead to performance evaluations that do not reflect real-world inference challenges in drug repurposing. To this end, we define three sets on which we evaluate KGE models: (i) a random test set, built by randomly sampling indication triples from the KG; (ii) a cold-start test set, also sampled from the KG, but constructed such that test drugs (or diseases) do not appear in any indication triples in the training data; and (iii) an independent test set, composed of curated drug-rare disease indications from the Orphanet database (13). Performance comparison assesses whether standard test constructions reflect realistic drug repurposing conditions.

## Material and Methods

### Definitions

#### Knowledge Graph Embedding and Link Prediction

A Knowledge Graph (KG) is a directed multi-relational graph representing entities and relationships as triples (*h, r, t*), where *h* and *t* are head and tail entities connected by relation *r*. Knowledge Graph Embedding (KGE) models encode these entities and relations into continuous vector spaces that capture latent KG patterns. Embeddings are learned by comparing positive and negative triples, with negative triples generated through Negative Sampling. A detailed formalization is available in Supplementary Section S1.

In link prediction, the goal is to complete missing triples by predicting either *t* in (*h, r*, ?) or *h* in (?, *r, t*). Each KGE model uses a scoring function to quantify triple plausibility. During evaluation, given *h* (or *t*) and *r*, all candidate entities are ranked by score, with the highest-ranked selected to complete the triple and infer missing KG links.

#### Training, Test, and Inference Sets

In this study, we define the following sets of triples:

- *Training Set:* The training set refers to triples sampled from the KG and used during the model training phase. In practice, this set is composed of two subsets: the training subset, which is directly used for optimizing the model parameters, and the validation subset, which is used to tune hyperparameters and prevent overfitting. For simplicity, we collectively refer to these subsets as the training set, unless the distinction is explicitly made.
- *Test Set:* The test set consists of triples sampled from the KG that were set aside and hidden during the training phase. For each triple in the test set, either the head or tail entity is masked, and the model’s ability to rank the correct entity among all possible entities is evaluated.
- *Inference Set:* In real-world deployment, an inference set would consist of incomplete triples in which either the head or tail entity is unknown and needs to be predicted. Here, we leverage an external dataset containing known drug-rare disease indications curated from Orphanet (Orphanet Drug-Rare Disease Indications). We use this dataset as an independent test set to evaluate the performance of the models in conditions that mimic real-world inference. We expect this analysis to reflect realistic drug repurposing applications for rare diseases.

### Data

#### Knowledge Graphs

**ShepherdKG** (14) is a biomedical KG dedicated to rare disease diagnosis. This KG was derived from PrimeKG (15), a KG developed for precision medicine applications, and includes additional knowledge from Orphanet (13), a comprehensive database for rare diseases. ShepherdKG contains 10 entity types, 30 relationship types and over 5 million triples (Supplementary Table S1).

**BioKG** (16; 17) is a biomedical KG developed for relational learning. It integrates information from multiple biomedical databases and contains 6 entity types, 17 relationship types and over 2 million triples (Supplementary Table S2).

**HetionetKG** (18) is a biomedical KG integrating data from public databases and designed to support drug repurposing through network-based inference. It contains 11 entity types, 24 relationship types and over 2 million triples (Supplementary Table S3).

All three KGs were divided as follows: 90% of the triples were assigned to the training set (including 10% of the triples reserved as the validation subset); the remaining 10% of the triples were allocated to the test set.

#### Orphanet Drug-Rare Disease Indications

To create the independent test set, we obtained access to Orphanet’s (13) dataset of Orphan drugs indicated for rare diseases, comprising 4915 drug-rare disease associations.

For each KG, we retained drug-rare disease associations involving drug and disease entities present in the KG to constitute the independent test set. Those that were already included in the KG were removed to ensure they were only present in the independent test set.

For ShepherdKG, 829 associations were retained as the independent test set. The majority of the excluded associations involved gene therapies, which are entities not included in ShepherdKG. Among the 829 retained drug-disease associations, 164 were already included in ShepherdKG and were removed from the KG to ensure they were only present in the independent test set. For BioKG, 292 associations were retained (69 were already included in BioKG and removed). For Hetionet, only 12 associations were retained (3 already included and removed).

### Models and Implementation

#### Models

We investigated two families of KGE models. The first family comprises decoder-only models (also referred to as shallow models), including the translational models TransE (19) and TransD (20) and the bilinear models RESCAL (21), DistMult (22), ANALOGY (23), ComplEx (24), and HolE (25).

Translational models represent relations as transformations in the embedding space, typically aiming to capture the relationship between entities through geometric operations like translations or rotations in the embedding space. In contrast, bilinear models use multiplicative interactions between embeddings, often through tensor operations (26).

The second family comprises Graph Neural Network (GNN)-based models (also referred to as deep models). These deep models have an encoder-decoder architecture. The GNN encoder computes entity embeddings by message passing, iteratively aggregating information from each entity’s neighborhood. A decoder then scores triples from the resulting embeddings. GNN-based models differ fundamentally from decoder-only models in how entity embeddings are learned. In a decoder-only model, each entity embedding is a free, independently learned parameter; in a GNN-based model, embeddings are computed from the training graph structure, so an entity’s representation depends on its neighbors’ embeddings.

We tested two GNN encoders, GAT (27) and GCN (28), each combined with a RESCAL decoder. For both encoders, we used a single message-passing layer, so that each entity’s embedding aggregates information from its direct (one-hop) neighbors only. Both encoders aggregate information at two levels: an intra-relation level, where a node’s neighbors’ representations are aggregated within each relation type, and an inter-relation level, where the resulting per-relation representations are combined across all relation types (using a sum aggregation in our experiments). The two GNN encoders differ in their intra-relation aggregation. In GAT, each neighbor’s contribution is weighted by a learned attention coefficient. In GCN, each neighbor’s contribution is instead combined using a fixed intra-relation aggregation function. We evaluated two such aggregation functions for our GCN encoders: mean aggregation (GCNmean) and max aggregation (GCNmax).

A detailed description of the models is provided in Supplementary Section S1. All decoder-only models were implemented using TorchKGE Python library (29).The GNN-based models were implemented using the KGATE Python library (30).

#### Negative Sampling

TorchKGE and KGATE implement three negative sampling strategies: *Uniform* (random entity replacement with uniform probability), *Bernoulli* (replacement probability based on relation mapping characteristics to minimize false negatives (31)), and *Positional* (replacement with entities previously observed at the same position for that relation, producing more realistic negatives (32)). For each positive triple, we generated five negatives using Uniform, five using Bernoulli, and one using Positional sampling.

#### Evaluation metrics

To evaluate the models on link prediction tasks on the validation set (i.e. the subset of the training set), test set, and independent test set, we used the filtered Mean Reciprocal Rank (MRR), a metric that measures the quality of ranked predictions by assessing the position of the first correct entity among all candidates ranked by the decoder. Higher MRR values indicate better performance, reflecting the model’s effectiveness in prioritizing correct links. Additionally, we occasionally report the filtered Hit@10 metric, which measures the proportion of correct entities ranked among the top 10 predictions. More details on these metrics are given in Supplementary Section S1.7.

#### Model Training

Models were trained using PyTorch (33) and PyTorch-Ignite (34) with Adam optimizer (learning rate 0.001, weight decay 0.001), embedding dimension 200, and batch size 4096. Translational models used Margin Loss (margin= 1); bilinear models used Binary Cross-Entropy Loss.

For decoder-only models, two training settings were used: (i) Standard training: validation every 20 epochs using MRR, early stopping after 10 consecutive non-improvements (maximum 500 epochs); (ii) Scheduled training: identical to standard training with added Cosine Annealing Warm Restarts scheduler (*T* 0 = 10, *T mult* = 2).

For GNN-based models, whose training is substantially more computationally demanding, we kept the same two settings but halved both the validation interval (to every 10 epochs) and the early stopping patience (to 5 consecutive non-improvements).

For all experiments, we reported the best MRR of the two trainings on the test set. Full MRR and Hit@10 results are available in Supplementary File S1.

#### Hardware and computational environment

All experiments were conducted on the Jean Zay supercomputer. For decoder-only models, we used a single NVIDIA V100 GPU with 32 GB of memory. For GNN-based models, we used a single NVIDIA A100 GPU with 80 GB of memory.

### DL1 - Redundancy between Training and Test Sets

In KGE-based link prediction tasks, redundancy between training and test sets due to inadequate data separation (called here Data Leakage 1, DL1) can arise from two distinct scenarios, as described by *Akrami et al*. (7).

First, when splitting the dataset into training and test sets, overlapping information between the splits may go undetected. This happens when the KG contains semantically related relations, known as near-duplicate or near-reverse-duplicate relations (7). For example, in a biomedical KG, relations like “(protein)-binds to-(protein)” and “(protein)-interacts with-(protein)” can be considered *near-duplicates* as they convey similar meanings. Similarly, reverse relations, such as “(drug)-inhibits-(protein)” and “(protein)-is inhibited by-(drug)” can be considered as *near-reverse-duplicates* as they convey symmetric semantic meanings. When these semantically related relations are randomly distributed across training and test sets without proper control, they create unwanted information leakage. Overall, failing to identify and account for semantically redundant relations can compromise the separation of the training and test data. As proposed by *Akrami et al*., semantically redundant relations can be detected by investigating whether, for two given relations, the heads and tails overlap exceeds predefined thresholds *θ*_1_ and *θ*_2_. These two thresholds account for asymmetric redundancy, where one relation may be redundant with respect to another without implying mutual redundancy (7). The complete mathematical formulation of this redundancy detection process is described in the Supplementary Section S1.8 (Equations S1 and S2).

Second, the KG may include *Cartesian product* relations, in which a large proportion of possible head-tail entity pairs are connected. In a biomedical KG, an example of such Cartesian product relation could be “(gene)-expressed in-(cell type)” for ubiquitously expressed genes (i.e. housekeeping genes). Cartesian product relationships can be identified by comparing the number of observed connections between the heads and tails covered by the relation with the total number of possible connections, and determining whether this proportion exceeds a predefined threshold *θ* (7). The specific mathematical formulation for this identification is provided in the Supplementary Section S1.9 (Equation S3).

#### DL1 Detection and Control Procedure

In order to detect and control DL1, we designed a preprocessing procedure to identify semantically redundant and Cartesian product relations within the KG, and ensure appropriate separation between training and test sets. This protocol is divided into several steps described below.

*Step 1: Filtering Reverse Triples* For every relationship type *r*, if a triple (*a, r, b*) exists in the KG, its reverse triple (*b, r, a*) is removed from the KG. This step addresses cases where the same relation *r* is used in both directions, which can be problematic since some KGE models (such as TransE and TransD) cannot encode symmetric relationships, while others (like DistMult) inherently impose symmetry on all relations. For undirected relationships, triple directionality is handled separately in Step 3 of this procedure.

*Step 2: Identifying Semantically Redundant and Cartesian Product Relations* For every possible pair of relations (*r*_1_, *r*_2_), we compute their semantic redundancy to identify near-duplicate and near-reverse-duplicate relations, using thresholds *θ*_1_ and *θ*_2_ set to 0.8. Additionally, for every relationship *r*, Cartesian product relations are identified using a threshold *θ* set to 0.8. Relations identified as semantically redundant are flagged for subsequent processing.

*Step 3: Handling Directionality for Undirected Relations* While most KGE models operate on directed triples, biomedical KGs often contain inherently undirected relationships. To accommodate these undirected interactions in our models, we systematically generate reverse triples with a new relation name. For any undirected relation *r*, we add a reverse counterpart *r*_rev_. Specifically, for every triple (*a, r, b*), we add (*b, r*_rev_, *a*). Because *r* and *r*_rev_ convey the same semantic information, the pair (*r, r*_rev_) is flagged and added to the list of semantically redundant relations identified in Step 2.

*Step 4: KG Split with Controlled Train-Test Separation* At this stage, semantically redundant relations and Cartesian product relations have been identified and flagged. The KG is split into training and test sets, with the following considerations:

- For semantically redundant relations flagged in Step 2, as well as relationships made directed by the addition of their reverse in Step 3, redundant triples are removed from the training set. Specifically, for any triple placed in the validation or test set, its redundant counterparts are excluded from the training set.
- For Cartesian product relations flagged in Step 2, we ensure that all triples involving a given head (or tail) with the Cartesian relation *r* are placed in the same split.

#### Evaluation of DL1

To evaluate the impact of DL1 on KGE model performance, we compared link prediction results obtained on the KGs with and without the application of the DL1 detection and control procedure described above. To create the dataset with DL1, steps 1 to 3 of the DL1 detection and control procedure were applied, but instead of using step 4’s controlled splitting procedure, the data was randomly split into training and test sets. To create the dataset without DL1, the full DL1 detection and control procedure was applied, including step 4, thereby ensuring that no redundant triples remained between the training and test sets.

For both datasets, KGE models were trained using the same hyperparameters and evaluation metrics for consistency. Link prediction performance was measured for the prediction of all relations (i.e., global performance), for the prediction of relations flagged in the DL1 detection and control procedure (i.e., performance on semantically redundant and Cartesian product relations), and for the prediction of the indication relation.

### DL2 - Investigating Node Degree as an Illegitimate Predictive Feature

To evaluate whether KGE models rely on node degree as an illegitimate feature for predicting new links (called here Data Leakage 2, DL2), we designed a permutation experiment. The underlying rationale is that if models utilize node degree as a predictive shortcut rather than learning meaningful semantic patterns, their performance should remain relatively stable when connections are randomized while preserving entity degrees. Formally, for a given relation *r*, we randomly permuted the tail entities while preserving the degrees of the heads and tails. Specifically, for a relation *r* connecting the set of heads *H* and the set of tails *T*, each tail in a triple is replaced by selecting a new entity randomly from the set *T*.

For this experiment, we used the KGs where DL1 had been controlled using the DL1 detection and control procedure described in DL1 Detection and Control Procedure. The permutation was applied to the indication relation between steps 3 and 4 of the DL1 detection and control pipeline. We compared the performance of KGE models trained on the permuted versus original version of the KG.

### DL3 - Sampling Bias in Test Set Construction

To investigate the presence and impact of sampling bias in test set construction (called here Data Leakage 3, DL3) in the context of drug repurposing, we compared KGE model performance on indication triples from the test set (i.e., triples sampled from the KG) and on an external independent test set of drug-rare disease indications (Orphanet Drug–Rare Disease Indications) reflecting realistic deployment scenarios. This comparison aimed to reveal the impact of dependencies that may exist between training and test sets, but are not expected between training and independent test sets. We conducted this analysis using two data splitting strategies: a standard random split and a cold-start split.

#### Standard Random Split

We used the same models as those employed for the DL1 experiment, trained on each KG after DL1 had been controlled through the procedure described in DL1 Detection and Control Procedure. In this experiment, we reported only model performance on predicting indication triples from the test set (i.e. indication triples sampled randomly from the original KG). We compared these performances with the performances obtained in the independent test set composed of curated drug-rare disease indications described in Orphanet Drug–Rare Disease Indications. This setup allowed us to evaluate whether models trained and evaluated on indication triples randomly sampled from each KG can generalize to predict drug-rare disease indications.

#### Cold-Start Split

We implemented cold-start splitting procedures applied to the indication relation in two distinct settings: a drug-based cold-start, and a disease-based cold-start. In the drug-based cold-start, all indication triples involving a given drug were assigned to the same split. Similarly, in the disease-based cold-start, all indication triples involving a given disease were assigned to the same split. These procedures ensured that models were required to predict indications for drugs or diseases that did not appear in any indication triples during training. Note that the cold-start splitting procedure is only applied to the indication triples. All drug and disease nodes are present in the training KG because of their other relations, hence their embeddings are learned during training.

To preserve a representative mix of frequent and rare entities while ensuring that each entity appeared in only one split, we stratified drugs or diseases into frequency bins based on their number of known indications and distributed them proportionally across the training, validation, and test sets. As shown in Supplementary Figures S1-S3, the resulting distributions were well balanced across splits, except for BioKG, where structural constraints of the graph led to a bias toward less-indicated entities in the validation and test sets (see Supplementary Figure S2 caption for details).

We then trained KGE models separately on the drug-based and disease-based cold-start splits, and evaluated each model on its corresponding test set and on the orphan drug independent test set. This allowed us to assess whether removing dependencies between training and test improved the alignment between test and independent test performance—i.e., whether cold-start splitting led to more realistic and reliable evaluation of generalization for drug repurposing tasks.

### Availability

The KGs used in this study are publicly available:

- ShepherdKG: https://zitniklab.hms.harvard.edu/projects/SHEPHERD/
- BioKG: https://github.com/dsi-bdi/biokg/releases/tag/v1.0.0
- HetionetKG: https://github.com/hetio/hetionet

The dataset of orphan drugs is available upon request from Orphanet at https://www.orphadata.com/samples/. All code and results are openly available on GitHub at https://github.com/galadrielbriere/data_leakage_kge_benchmark.git. The repository provides both the implementation to reproduce the results obtained in this study and a reusable framework for assessing data leakage in any KG, including the DL1 detection and control procedure, the DL2 permutation procedure, the DL3 cold-start split procedure, and the complete models and training implementations.

## Results

Experiment 1: Assessing the Impact of Redundancy between Training and Test Sets (DL1) on Model Performance

Existing benchmarks on biomedical KGs using similar KGE models and reporting MRR values show MRR ranging from 0.1 (12) to 0.8 (9), and even up to 0.9 (35). This large range in performance may stem from differences in KG structure and model training protocols. However, DL1 (i.e., presence of redundancy between training and test data) has not been systematically investigated in these benchmarks, leaving open the question of whether data leakage contributes to these observed performance disparities.

To assess the impact of DL1 on model performance, we trained and evaluated seven decoder-only (TransD, TransE, DistMult, RESCAL, HolE, ComplEx and ANALOGY) and three GNN-based (GCNmax, GCNmean and GAT, all combined with a RESCAL decoder) KGE models (Material and Methods) on two variants of ShepherdKG: the ShepherdKG where only the steps 1 and 3 of the DL1 detection and control procedure (Material and Methods) were applied, followed by random train-test splitting (i.e., KG with DL1), and the ShepherdKG where all four steps of the DL1 detection and control procedure were applied to remove any redundancy between training and test sets (i.e., KG without DL1).

The result of the application of the DL1 detection and control procedure on the ShepherdKG resulted in the removal of 1, 336, 637 reverse triples at Step 1. Most of these removed triples (1, 336, 314) were drug-drug relations, as for every (*h*, drug-drug, *t*), the reverse triple (*t*, drug-drug, *h*) was also included in the KG. At Step 2, we did not identify any semantically redundant or Cartesian product relations (using *θ*_1_, *θ*_2_, and *θ* set to 0.8). Because most KGE models consider directed relations, three undirected relations were made directed at Step 3: drug-drug similarity (*drug drug* relation), disease-disease similarity (*disease disease* relation), and protein-protein interactions (*protein protein* relation). For these relations, the reverse was included with a different relation name (e.g., *drug drug rev* ) to avoid introducing symmetric relationships. Including these reverse triples created a semantic redundancy in the KG, which was removed at Step 4 of the DL1 detection and control procedure for the KG controlling for data leakage but retained for the KG where data leakage was not controlled. Our results revealed that redundancies between training and test data could substantially inflate model performance, as measured by global MRR (Figure 1-a). However, the extent of this inflation varied across models and may depend both on the intrinsic capacity of the model and its ability to exploit the leakage.

**Figure 1.**
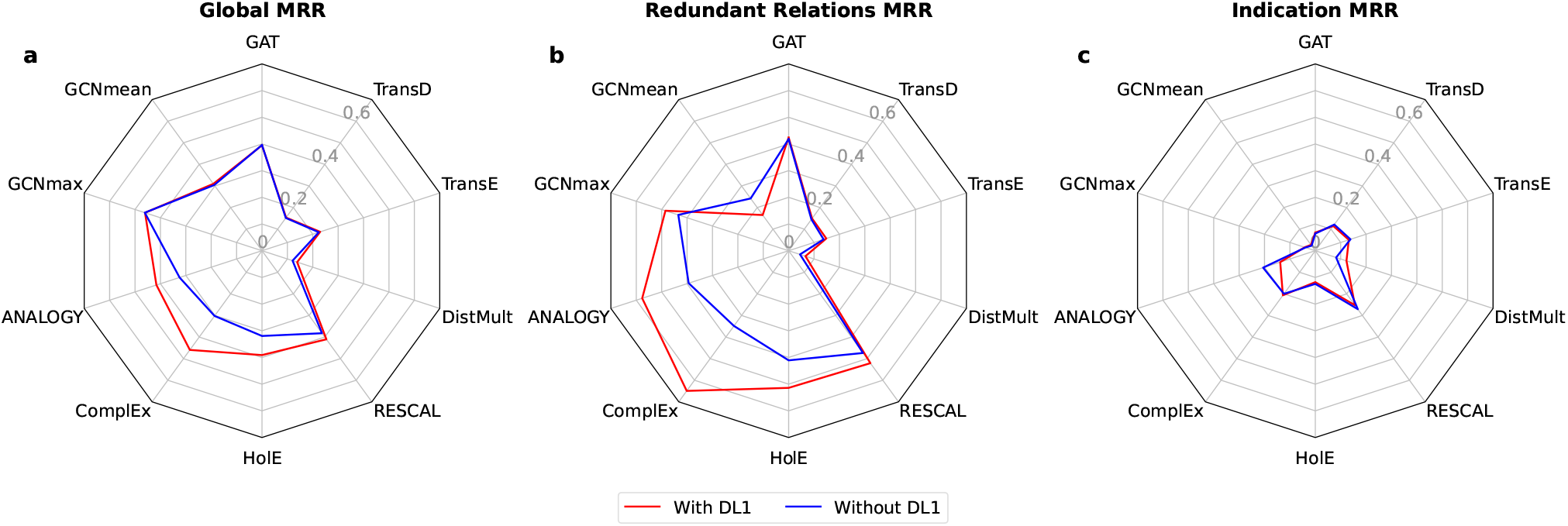
Impact of Training-Test Set Separation on KGE Model Performance (DL1) on ShepherdKG. **(a)** Global MRR: performance of the models across all relations in the test set. **(b)** MRR for Redundant Relations: Model performance on the relations flagged as redundant by the DL1 detection and control procedure. **(c)** MRR for the Indication Relation: Model performance on indication triples. Performance shown with (red) and without (blue) DL1

Among the best performing decoder-only models, ComplEx showed the most pronounced impact of DL1: the global MRR dropped from 0.46 to 0.30 (a 34.7% decrease), when DL1 was removed. ANALOGY and HolE also exhibited performance decreases when DL1 was controlled (dropping from 0.41 to 0.32 and 0.39 to 0.32 respectively). In contrast, RESCAL maintained high performance regardless of DL1 (with an MRR dropping from 0.41 to 0.38, making it the best-performing decoder-only model when DL1 was removed). TransE also showed only marginal difference (from 0.23 to 0.22) but with lower overall performance. Finally, DistMult and TransD both exhibited low overall performance and minimal change when DL1 was controlled.

The GNN-based models showed overall good performance and were essentially unaffected by DL1 control, their global MRR changing by at most 0.006 between the with-DL1 and without-DL1 settings. GCNmax and GAT were the best-performing models in the absence of DL1, with global MRRs of 0.46 and 0.40 respectively.

In addition to analyzing the global MRR, which considers all the KG’s relations, we computed the MRR for relations that were semantically redundant between the training and test sets. These semantically redundant relations are causing DL1 (Figure 1-b). For these relations, the gap between the reported performance of decoder-only models trained with and without DL1 was even more pronounced, as expected. For ComplEx, ANALOGY and HolE, the MRR decreased by 46%, 32%, and 20%, respectively, when DL1 was removed. RESCAL, however, continued to report nearly equal MRR values across the two settings (decreasing from 0.52 to 0.47, and remaining the best performing model when DL1 is controlled), similar to its performance for the global MRR. DistMult, TransE, and TransD showed poor performances, regardless of the presence of DL1, with MRR values below 0.15. However, considering separately the three types of semantically redundant relations nuanced this picture (Supplementary Figure S4): while DistMult, TransE, and TransD models showed no meaningful difference for the prediction of drug-drug and protein-protein interactions upon DL1 removal, they did exhibit lower performance on disease-disease relations. This indicates that those models are also affected by DL1 at the relation-specific level. Strikingly, for disease-disease relations, models such as ComplEx and ANALOGY reached MRR values of up to 0.6 in the presence of DL1, yet dropped below 0.1 when DL1 was removed. This collapse reveals that the apparent correct predictive performance on this relationship type is entirely driven by DL1 for these decoder-only models.

On semantically redundant relations, GNN-based models varied in performance. GAT and GCNmax reported comparable or slightly higher MRR with DL1 than without (GAT remaining nearly unchanged at 0.42 and GCNmax increasing from 0.43 without DL1 to 0.49 with DL1). GCNmean, in contrast, performed worse with DL1 (0.17) than without (0.24). GCNmean was the only model for which controlling DL1 improved performance rather than degrading it. We hypothesize that this behavior in the GCNmean model arises from the interaction between mean aggregation and graph density. Mean aggregation averages a node’s neighbors’ representations within each relation type, so that each neighbor contributes only a fraction 1*/k* of the aggregated signal, where *k* is the number of neighbors. By retaining redundant counterpart triples in the training set, DL1 increases graph density and enlarges node neighborhoods, diluting the contribution of any individual neighbor and the useful structural signal. Max aggregation used in the GCNmax model, in contrast, does not average over the node’s whole neighborhood, so strong signals are preserved rather than diluted. This may make GCNmax robust to graph densification, which is consistent with GCNmax outperforming GCNmean. Considering separately the three types of semantically redundant relations (Supplementary Figure S4) supports this hypothesis: with DL1, GNN-based models reached higher or comparable MRR than without DL1 across the three redundant relation types. The only case where this pattern was reversed was for GCNmean on drug-drug interactions. This fits with the interaction between mean aggregation and density described above described above: including DL1 adds far more edges to the drug-drug relation than to any other relation (about 214k added triples in the training ShepherdKG, as compared to about 51k for proteinprotein and 5k for disease-disease relations), so this is where mean aggregation most dilutes each neighbor’s contribution.

Overall, the results clearly indicate that KGE models are sensitive to DL1, and our findings highlight the significant impact of DL1 on the evaluation of KGE models, particularly for semantically redundant relations, where its presence can lead to artificially high evaluation metrics. These findings underscore the critical importance of controlling DL1 during model evaluation.The inflation in performance due to DL1 affects not only decoder-only models but also, to a lesser extent, GNN-based ones. This raises the question of how such inflation arises, and why its magnitude differs between decoder-only and GNN-based models. We address this in Supplementary Section S2.3, where we describe the possible mechanisms through which DL1 can impact performance: a decoder-level channel, common to all models, and an additional message-passing channel specific to GNN-based models.

In addition, we wondered whether DL1-affected relations could potentially help the prediction of non-DL1 affected relations. Specifically, the indication relation was not semantically redundant in the KG (i.e. not affected by DL1). We tested if the presence of DL1-affected relations in the KG might, in some way, assist in predicting the non-DL1 affected indication triples. The results in Figure 1-c were inconclusive. The prediction performance was overall low in all models, reaching 0.27 at best (RESCAL model in DL1 controlled setting). These results suggest that introducing semantic redundancy through DL1, in addition to compromising the reliability of model evaluation, does not provide any consistent benefits for predicting non-redundant relations like indication.

To verify that these findings generalized beyond ShepherdKG, we replicated the full analysis on two additional biomedical KGs: BioKG and HetionetKG (Supplementary Section S2.2). The three patterns reported above held across all KGs: (i) global MRR was inflated by DL1, most strongly for the decoder-only models, while GNN-based models were less affected (Supplementary Figures S5 and S6); (ii) this inflation was even more pronounced on semantically redundant relations (Supplementary Figures S7 and S8); and (iii) DL1-affected relations did not consistently help predict non-redundant relations such as drug-disease indications (Supplementary Figures S5 and S6).

### Experiment 2: Investigating Node Degree as an Illegitimate Predictive Feature (DL2)

A second potential source of data leakage we examined was the use of node degree as an illegitimate feature. To test this data leakage source in the context of indication prediction, we designed an experiment in which the indication triples in the KG were permuted while preserving the original degree of entities (Material and Methods). Our hypothesis was that if the models rely solely on node degree as an illegitimate feature, the MRR for the indication relation should remain similar between models trained on the original and on the permuted KGs.

Our results on the ShepherdKG, presented in Figure 2-a, indicated that the decoder-only models did not rely solely on node degree as a predictive feature. Indeed, decoder-only models trained on the KG with permuted indication triples exhibited significantly poorer predictive performance for this relation as compared to the same models trained on the original KG (e.g., from 0.27 on the original graph to 0.02 on the permuted one for RESCAL, and from 0.21 and 0.20 to 0.02 for ANALOGY and ComplEx). Because degree remains available after permutation, this drop shows that the decoder-only models predict indications from the specific drug-disease associations rather than from degree alone. We further analyzed the geometry of the embeddings (Supplementary Section S2.4) to assess whether degree was encoded in the norms or direction of the entity embeddings. On ShepherdKG, degree was at best partially and unevenly encoded: drug degree is not reflected in the embedding norm or direction for any decoder-only model, while disease degree carried only a moderate norm signal for ANALOGY and ComplEx, and was weakly encoded in the direction of the entity embeddings for some models. The strength of this degree encoding did not drive predictive performance: the performance of several models was largely degraded even though disease degree was better encoded on the permuted graph than on the original one. Degree encoding thus explains neither the models’ accuracy on the original ShepherdKG nor the gap between the original and permuted graphs.

**Figure 2.**
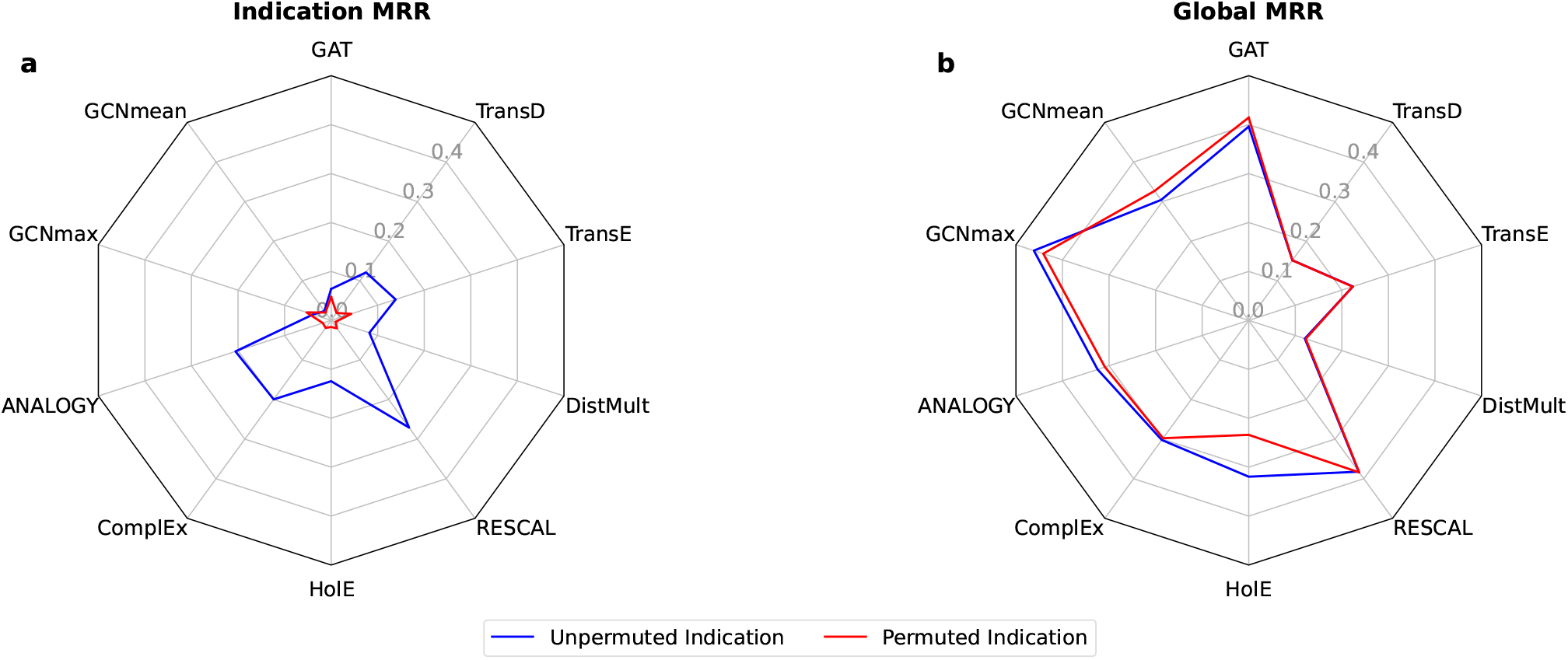
Assessment of node degree exploitation as an illegitimate feature (DL2) in indication prediction on ShepherdKG. **(a)** MRR for the Indication Relation: Model performance on indication triples. **(b)** Global MRR: Model performance across all relations in the test set. Performance shown with original indications triples (blue) and degree preserving permutation for indication triples (red).

However, permuting the indication triples did not affect model performance on predicting other types of relations, as the global MRR was similar for most models (Figure 2-b). This suggests that the indication relation does not provide information used by the models to predict other types of relations.

On BioKG (Supplementary Figure S9-a), the decoder-only models were poor predictors of the indication relation whether or not its triples were permuted (MRR ≤ 0.07). On HetionetKG (Supplementary Figure S10-a), the permutation degraded the decoder-only models only partially, without the clear collapse observed on ShepherdKG. We however interpret this result with caution as the indication test set on HetionetKG contains only 76 triples, compared with 844 on ShepherdKG and 6,611 on BioKG; on a set this small, the estimated MRR may not be reliable.

The GNN-based models showed more mixed behavior. On ShepherdKG, they predicted the indication relation poorly, whether or not its triples were permuted (all MRRs below 0.07, essentially unchanged by the permutation), leaving the DL2 experiment inconclusive on this KG (Figure 2-a). On BioKG, however, the GNN encoders retained substantial predictive performance on the permuted graph (Supplementary Figure S9-a). This was more striking for GCNmax, whose MRR was only marginally reduced by the permutation (0.54 MRR on the original graph, 0.50 on the permuted one). GAT was more sensitive to the permutation but still reached an MRR of 0.22 on the permuted graph. We showed that degree is encoded in the GNN representations (Supplementary Section S2.4), and since our permutation strategy preserves node degree, this information remains available to these models. However, the retained performance on BioKG may not be driven by node degree alone, since the permutation might have also preserved other topological properties accessible through message passing.

### Evaluating Sampling Bias in Test-Set Construction and Its Effect on Evaluation Reliability (DL3)

The reliability and utility of the performance metrics obtained on the test set depends on how well the test set correctly represents real-world inference scenarios. In standard practice, test sets are randomly sampled from the same KG that is used for training. As a result, test and training sets often share similar statistical and structural properties. However, at deployment time, the models might encounter entities and relationships with substantially different characteristics compared to those observed during training and evaluation. These differences can potentially lead to overoptimistic estimates of model performance.

In this experiment, we investigated whether performance on such randomly sampled test sets reliably predicted performance on an independent test set composed of drug-rare disease indications from Orphanet (13) (Orphanet Drug-Rare Disease Indications). In addition to the random split, we further considered a cold-start splitting strategy, where all known indications for a given drug or disease were assigned to the same training or test split. The resulting cold-start test split is expected to better reflect realistic drug repurposing scenarios, where models are expected to make predictions for understudied drugs or diseases with few or no known indications in the training data.

#### Random Split

We first examined whether evaluating models on test data randomly sampled from the same KG as the training data (Standard Random Split) provided reliable performance indicators for real-world drug repurposing inference. To do so, we compared the performance of the models obtained on the test set and on our independent test set of drug-rare disease indications (Orphanet Drug-Rare Disease Indications).

For ShepherdKG, the results, shown in Figure 3-a, indicated that while the best-performing decoder-only models achieved MRRs around 0.2 on the test set (RESCAL, ComplEx, ANALOGY), their performance on the independent test set was close to 0. The best-performing decoder-only model on the independent test set, RESCAL, achieved an MRR of only 0.016, with a Hit@10 of 0.031, meaning that, on average, the correct entity was ranked among the top 10 predictions in just 3.1% of the cases. In comparison, the same model achieved a Hit@10 of 0.46 on the test set indications, successfully ranking the correct entity in the top 10 predictions in nearly half of the cases. This gap between test set and independent test set performance was also observed on BioKG and HetionetKG (Supplementary Figures S11-a and S12-a).

**Figure 3.**
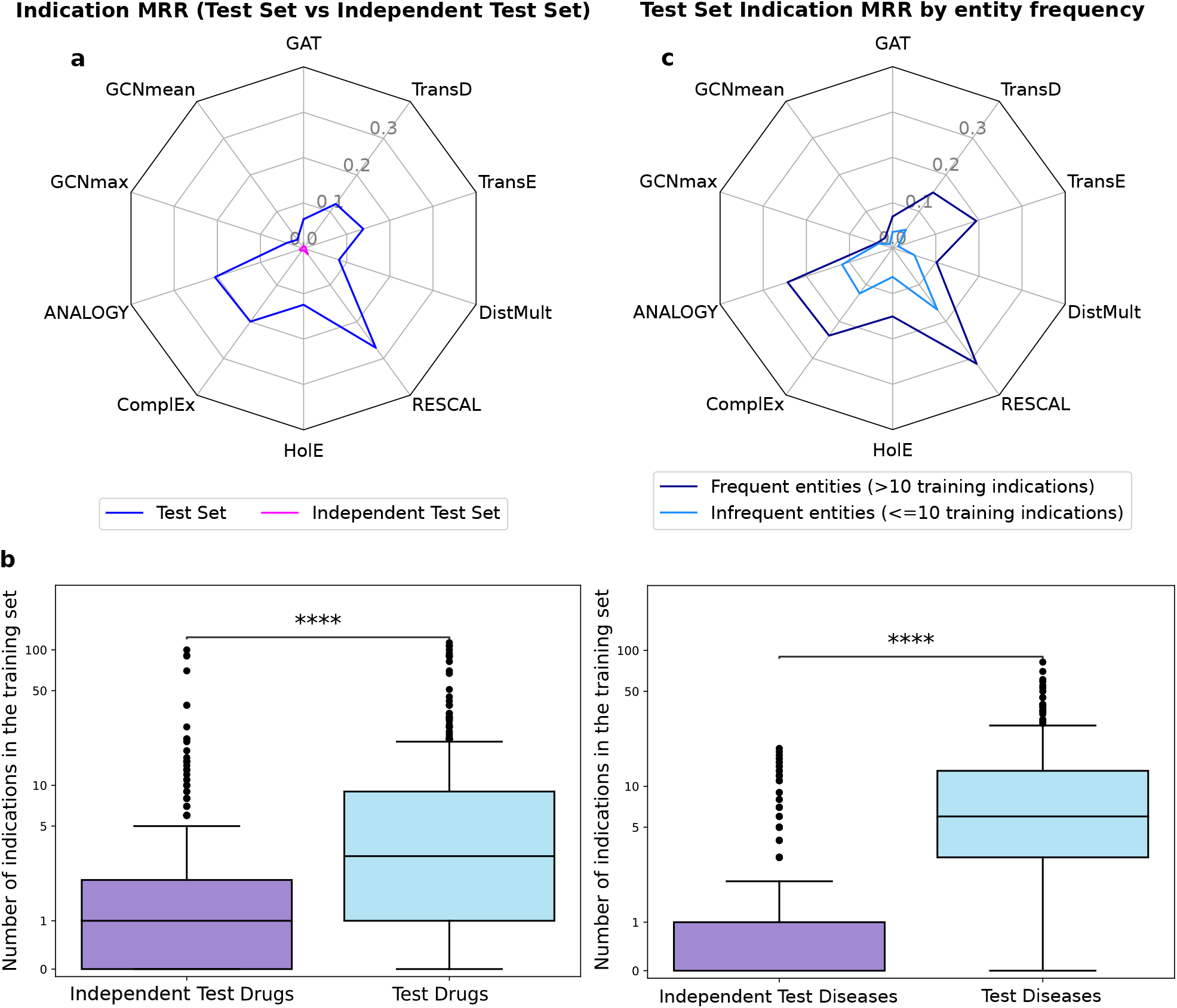
Random Split Test Set and Independent Test Set Comparisons (DL3) on ShepherdKG. **(a)** MRR comparison between test set of randomly sampled indication triples (blue) and independent test set indication triples (pink). **(b)** Distribution of the number of training set indications (shown on a logarithmic scale) per drugs and per diseases across test and independent test sets. Statistical significance was assessed using the Mann-Whitney-Wilcoxon test (two-sided). Annotations indicate significance levels: **** (*P* ≤ 1 × 10−4). **(c)** MRR comparison on test *indication* triples involving entities with frequent (*>* 10) or infrequent (≤ 10) occurrences in training *indication* triples (dark and light blue, respectively).

On ShepherdKG, the GNN-based models were already poor predictors of the indication relation on the test set, but we also observed a decrease of performance on the independent test set. On BioKG and HetionetKG, GNN-based models were among the best predictors of the indication relation on the test set. However, their performance also collapsed on the independent test set (Supplementary Figures S11-a and S12-a). On BioKG, GCNmax reached the highest test-set MRR of all models (0.54) yet dropped to 0.002 on the independent test set, while GCNmean and GAT fell from 0.20 and 0.43 to 0.003 and 0.006, respectively. The same collapse was observed on HetionetKG, where GCNmax, GCNmean and GAT dropped from 0.25, 0.24 and 0.33 on the test set to 0.03, 0.01 and 0.01 on the independent test set.

To understand the reasons behind such poor generalization on the independent test dataset, we analyzed the connectivity patterns of drug and disease entities. For each entity belonging to either the test or the independent test set, we computed the number of indication triples it participates in in the training set. In ShepherdKG, we found that drugs and diseases from the test set had significantly higher numbers of known indications in the training set as compared to drugs and diseases from the independent test set (Figure 3-b). Additionally, we computed the total degree of each entity in the training set—that is, the number of triples it was involved in across all relation types (Supplementary Figure S13). Drugs in the independent test set had a lower median total degree (529) compared to drugs in the test set (1196), and diseases in the independent test set were also slightly less connected (median degree of 33 vs. 41 for diseases in the test set). These results confirmed that entities from the independent test set were globally less connected in the training ShepherdKG, which may partly explain the models’ poor generalization ability. The same pattern was observed in BioKG and HetionetKG (Supplementary Figures 14 and 15).

We further investigated whether test triples involving drugs or diseases with more frequent indications in the training set were predicted more accurately than those involving rarer ones. To do so, we selected indication triples from the test set and split them into two groups: triples in which either the drug or the disease appeared in more than 10 indication triples in the training set (frequent group, 590 triples), and triples where both entities had 10 or fewer indication triples (infrequent group, 254 triples). As shown in Figure 3-c, all models performed significantly better on test triples involving entities (drugs or diseases) that appeared in more indication triples in the training set. Similar patterns were observed for BioKG and HetionetKG (Supplementary Figures 11-c and 12-c).

These experimental results demonstrate significant limitations of current KGE models for drug repositioning applications. While these models achieve reasonable performance on the test set, their predictive capabilities exhibit a strong dependence on the number of known indications in the training data involving the specific drug or disease under consideration. The models systematically underperform when predicting indications for drugs or diseases with few known indications and show particularly poor generalization to rare disease applications. The reliance on a high number of known indications severely limits the models’ practical applicability, as the identification of novel indications for less-studied drugs and diseases represents a critical need.

#### Cold-start Splits

To assess whether test set construction can better reflect the challenges of real-world drug repurposing inference, we designed a cold-start splitting protocol that we applied on the indication relation, as described in Cold-Start Split. We implemented two variants of this protocol: a drug-based cold-start, where all indication triples involving a given drug were assigned to a single split, and a disease-based cold-start, where the same constraint was applied to diseases. This setup ensured that test entities had no associated indications in the training set, thus mimicking inference conditions for novel or understudied drugs or diseases.

For ShepherdKG, as shown in Figure 4-a, models achieved higher performance on the test set under the random split compared to both cold-start configurations, which was expected since the random setting allows drugs and diseases to be partially known through their training indications. In contrast, the cold-start settings presented a more challenging prediction task. For ShepherdKG, we observed a substantial and consistent drop in test performance when moving from the random to the cold-start splits. The two best-performing models on the random split, RESCAL and ANALOGY, achieved MRRs of 0.27 and 0.21, respectively. These values dropped to 0.16 and 0.09 in the drug-based cold-start setup, and further down to 0.09 and 0.07 in the disease-based cold-start, where the task proved even more difficult. Similar trends were observed for all other models, confirming that the performance degradation were not model-specific but reflected a general limitation of current KGE approaches. This consistent drop indicates that KGE models rely heavily entity-specific exposure to the indication relation during training to make accurate predictions, and struggle to generalize when drugs or diseases lack such exposure despite appearing in other contexts.

**Figure 4.**
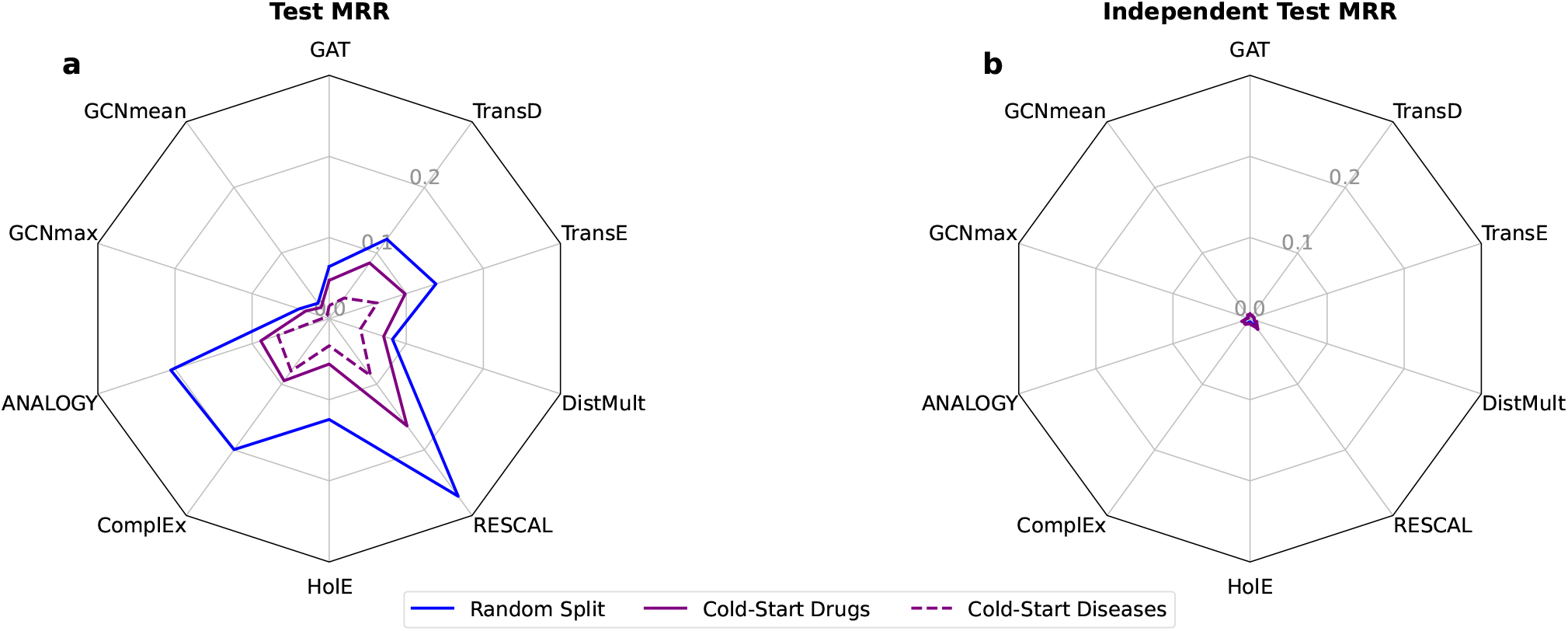
Evaluating Test Set and Independent Test Set Performance under Random vs. Cold-Start Splits (DL3) on ShepherdKG. **(a)** MRR comparison on the test set: randomly sampled indication triples (blue), drug-based cold-start indication triples (solid dark purple) and disease-based cold-start indication triples (dashed dark purple). **(b)** MRR comparison on the independent test set: randomly sampled indication triples (blue), drug-based cold-start indication triples (solid dark purple) and disease-based cold-start indication triples (dashed dark purple).

Nevertheless, models still retained some predictive capacity in the cold-start scenario. RESCAL, for instance, reached an MRR of 0.16 (hit@10 = 0.29) in the test set of the drug-based cold-start split. This suggests that even in the absence of indication triples for the considered drugs, structural signals from other parts of the graph remain informative. Still, the persistent performance gap between random and cold-start test sets emphasizes the difficulty of generalizing to new drug or new disease indications.

Similar patterns were observed on BioKG (Supplementary Figure S16-a). On Hetionet, the results were more mixed, with some models achieving higher test MRR under the drug- or disease-based cold-start splits, while others performing better under the random split (Supplementary Figure S17-a).

Figure 4-b shows model performance on our independent test set in ShepherdKG. The independent test performance remained low regardless of the test set sampling strategy. The best model in all settings, RESCAL, achieved at most an MRR of 0.016 and hit@10 of 0.03 (in the disease-based cold-start), comparable to the scores observed using random sampling. These results suggest that even though cold-start split offers a more realistic and challenging evaluation framework, it is not sufficient to bridge the gap between test performance and real-world inference capability. This pattern was confirmed in BioKG and HetionetKG (Supplementary Figures S16-b and S17-b).

Taken together, these findings call into question the reliability of standard test set construction strategies, such as random or cold-start sampling, for estimating the real-world inference capabilities of KGE models in drug repurposing scenarios.

## Discussion

Data leakage critically undermines the evaluation of KGE models by inflating performance metrics and obscuring real-world applicability. We systematically assessed the impact of three leakage sources on link prediction in a biomedical KG, showing the need to identify and mitigate these issues to obtain realistic performance estimates. Because KGE models remain largely black boxes, transparent reporting of leakage controls is essential for reliable and reproducible evaluations. We therefore advocate for standardized practices to detect and mitigate data leakage in KGE research and, more broadly, in machine learning.

In our first experiment, we showed that redundancy between training and test sets (DL1) can substantially inflate performance metrics, obscuring the true capabilities of KGE models. This highlights the need for consistent and transparent evaluation protocols. We developed a systematic procedure to detect and control DL1 in link prediction tasks and make it publicly available to support robust evaluation practices. We also identified challenges specific to undirected relations in KGs. Converting undirected relations into directed ones by adding reverse edges, often required because most KGE models operate on directed graphs, can itself introduce DL1. While the cleaning procedure we propose mitigates redundancy arising from reverse relations, the optimal treatment of undirected relations remains open. Further work is needed to determine whether using only the original direction or including reverse relations with carefully controlled redundancy yields better performance.

A striking finding from our DL1 analyses is that certain relation types appear harder to learn than others, independently of the presence of data leakage. Even among comparable relationship types such as drug-drug, disease-disease, and protein-protein interactions, which are all undirected and homogeneous, the same models achieve different performance levels in the absence of DL1. Understanding what drives these differences would require dedicated metrics that could explain relation-specific learnability.

In our second experiment, which examined whether KGE models rely on node degree as an illegitimate feature (DL2), we found no evidence that predictions were driven by degree alone. For decoder-only models, permuting the indication triples (while preserving degree), caused predictive performance to collapse, showing that degree information alone does not support indication prediction. The GNN encoders retained substantial performance on the permuted BioKG, but this may not be driven by degree alone either: the permutation might preserve not only the degree distribution but other structural features that message passing could exploit. These results suggest that degree-based permutation alone may not capture all forms of DL2, and that other illegitimate features may remain undetected. This is consistent with a recent study on drug synergy prediction, in which degree-preserving network rewiring similarly failed to reveal degree-based shortcuts (**?** ). In that study, however, the authors additionally permuted edge weights and found that the models were exploiting the aggregated weight of individual nodes as a predictive shortcut rather than meaningful pairwise interactions. As we do not have edge weights in our KGs, we cannot test this hypothesis directly. One promising direction for further investigation is the influence of community structure that may enable prediction shortcuts by emphasizing local connectivity over meaningful semantic relationships. Assessing how strongly KGE models are biased by such structural features is an important next step toward understanding their true capabilities.

In our third experiment, we showed that models performing well on a randomly sampled test set failed to generalize to an external real-world drug-repurposing dataset focused on rare diseases. This is the most concerning result of our study. This generalization gap was reinforced by stratifying performance by drug and disease indication frequencies: models performed markedly better for entities with more than ten known indications. This frequency-dependent pattern suggests that predictions rely heavily on local structural cues and direct exposure during training rather than on representations that support broader generalization.

To assess whether test set construction contributes to this performance mismatch, we implemented cold-start evaluation protocols that prevent test drugs or diseases from appearing during training for the target relation. Although performance decreased relative to the random split, a substantial gap between test and independent test performance persisted. This suggests that current evaluation strategies do not adequately reflect the generalization challenges involved in real-world drug repurposing inference.

Altogether, our results raise concerns about the suitability of current KGE methods for link prediction tasks, particularly in the context of drug repurposing. Even when evaluated under strict and realistic conditions, controlled for data leakage, models show a gap between test set performance and independent test set performance. This reveals a generalization issue: performance on withheld test data fails to transfer to independent evaluation, and ultimately to real-world inference. We argue that the field needs to move beyond benchmarking on withheld test datasets extracted from the training data, and toward evaluation frameworks that are more aligned with real-world inference scenarios. We see two promising directions. First, temporally stratified splits, where training data chronologically precedes test data, would allow prospective prediction tasks. Second, community-organized benchmarks and challenges could provide standardized, independently curated evaluation sets that more faithfully represent real-world inference challenges.

## Author contributions statement

G.B. and A.B. conceived the study. G.B. implemented, conducted the experiments, and analyzed the results. T.S. developed the proof of concept for the study. B.L. reviewed and contributed to the implementation. A.B. supervised the project.

G.B. and A.B. wrote the manuscript. All authors reviewed and approved the final manuscript.

## Acknowledgments

This work was supported by the French government under the France 2030 investment plan, as part of the Initiative d’Excellence d’Aix-Marseille Université - A*MIDEX (AMX-19-IET-007) through the Marseille Maladies Rares Institute (MarMaRa); the French National Research Agency through the CAMUDI project [ANR-21-CE45-0001]; and the European Rare Diseases Research Alliance (ERDERA, European Union’s Horizon Europe research and innovation programme under grant agreement [N°101156595]).This project was provided with computer and storage resources by GENCI at IDRIS thanks to the grant 2024-AD010315786 on the supercomputer Jean Zay’s V100 and A100 partition. We acknowledge Orphanet and INSERM for providing the dataset of orphan drugs used in this study. We thank Elias Ventre for his thorough manuscript review and valuable scientific input. We thank Léo Guignard for providing access to GPU resources used for development and debugging. We thank Antoine Toffano for his insightful feedback on the data leakage mechanisms. We used large language models (LLMs) to assist in coding and in polishing the writing of the manuscript.

