## Supplementary Materials for "Benchmarking the Impact of Data Leakage on the Performance of Knowledge Graph Embedding Models for Biomedical Link Prediction"

### Supplementary Material

Galadriel Brière,[1,*] Thomas Stosskopf,[1,2] Benjamin Loire[1,3] and Anaïs Baudot[1,4]

[1]Aix Marseille Univ, INSERM, MMG, Marseille, France, [2]Present address: Aix Marseille Univ, INSERM, TAGC, Marseille, France, [3]Neurology Therapeutic Area, R&D Servier Paris-Saclay Institut, Gif-sur-Yvette, France and [4]CNRS, Marseille, France

#### Contents

S1 Definitions and notations 1
S1.1 Knowledge Graph 1
S1.2 Knowledge Graph Embeddings 1
S1.3 Embedding Framework 2
S1.4 Encoder used in this study 2
S1.5 Decoders used in this study 3
S1.6 Negative Sampling 3
S1.7 Evaluation Metrics 3
S1.8 Semantically redundant relations 4
S1.9 Cartesian product relations 4
S2 Supplementary Results 4
S2.1 Sensitivity Analysis on $\theta_1$, $\theta_2$ and $\theta$ Thresholds 4
S2.2 DL1 detection and control procedure on BioKG and HetionetKG 4
S2.3 DL1 leakage operates through different channels in decoder-only and GNN models 5
S2.4 Degree can be encoded in entity embeddings without driving link prediction performance 6
S3 Supplementary Tables 7
S4 Supplementary Figures 8

#### S1. Definitions and notations

##### S1.1. Knowledge Graph

We define Knowledge Graphs as directed multi-relational graphs, formally represented by a tensor $\mathcal{A} \in \mathbb{R}^{|V|\times|V|\times|R|}$, where $V$ denotes the set of entities, $R$ represents the set of relations, and $\mathcal{A}_{i,j,r}$ indicates the presence of a directed interaction of type $r \in R$ from entity $v_i \in V$ to entity $v_j \in V$.

When entities are associated with specific node types, the KG can be further extended by including a type function $T : V \to C$, mapping each entity to one or more types from a set $C$ of possible types.

Additionally, if the entities are associated with additional features, such as numerical attributes, text, or images, the Knowledge Graph can be extended to a *multimodal* Knowledge Graph. Entity features are typically encoded into real-valued vectors (e.g., through pre-trained embeddings for text, images, or other types of data) and are represented by a feature matrix $X \in \mathbb{R}^{|V|\times d_0}$, where each row corresponds to the feature vector of an entity, and $d_0$ is the dimensionality of the encoded feature space. If no entity features are available, $X$ can be omitted.

Each interaction in the graph can be represented by a triple $(h, r, t) \in V \times R \times V$, where $h \in V$ represents the head entity, $t \in V$ the tail entity, and $r \in R$ the directed relation connecting $h$ to $t$.

This interaction is encoded in the tensor $\mathcal{A}$ such that:

$$\mathcal{A}_{h,t,r} = \begin{cases} 1, & \text{if there is an interaction of type } r \text{ from } h \text{ to } t, \\ 0, & \text{otherwise.} \end{cases}$$

##### S1.2. Knowledge Graph Embeddings

In KG representation learning, both entities and relations are typically embedded into continuous vector spaces. The embeddings of entities are collectively represented by the entity embedding matrix $\mathbf{Z} \in \mathbb{R}^{|V|\times d_e}$, where each row $\mathbf{z}_i \in \mathbb{R}^{d_e}$ corresponds to the embedding of entity $v_i \in V$. Similarly, the embeddings of relations are represented by the relation embedding matrix $\mathbf{R} \in \mathbb{R}^{|R|\times d_r}$, where each row $\mathbf{r}_k \in \mathbb{R}^{d}_{r}$ corresponds to the embedding of relation $r_k \in R$.

For a specific triple $(h, r, t)$, we denote the embeddings as $(\mathbf{h}, \mathbf{r}, \mathbf{t})$, where $\mathbf{h}, \mathbf{r}, \mathbf{t} \in \mathbb{R}^d$ are the respective embeddings of the head entity, relation, and tail entity in the vector space.

Thus, the embeddings $\mathbf{h}, \mathbf{r}, \mathbf{t}$ are rows of the entity and relation embedding matrices $\mathbf{Z}$ and $\mathbf{R}$, such that:

$$\mathbf{h} = \mathbf{Z}[h, :], \quad \mathbf{r} = \mathbf{R}[r, :], \quad \mathbf{t} = \mathbf{Z}[t, :].$$

#### S1.3. Embedding Framework

We extend the encoder-decoder framework introduced by Chami et al. (1), originally designed for network embedding, to support Knowledge Graphs.

This framework considers two key components; an encoder and a decoder, described as follows:

- **Knowledge Graph encoder** $\text{ENC}_{\Theta_{Enc}}$:

$$\text{ENC}_{\Theta_{Enc}} : \mathbb{R}^{|V|\times|V|\times|R|} \times \mathbb{R}^{|V|\times d_0} \rightarrow \mathbb{R}^{|V|\times d_e} \times \mathbb{R}^{|R|\times d_r}$$

The encoder is parameterized by $\Theta_{Enc}$ and combines the graph structure, represented as a tensor $\mathcal{A}$, with optional node features $X \in \mathbb{R}^{|V|\times d_0}$, where $d_0$ represents the dimensionality of the node features.

The encoder outputs both the entity embedding matrix $\mathbf{Z} \in \mathbb{R}^{|V|\times d_e}$ and the relation embedding matrix $\mathbf{R} \in \mathbb{R}^{|R|\times d_r}$, where $d_e$ and $d_r$ represent the embedding dimensions for entities and relations, respectively.

Formally:

$$\mathbf{Z}, \mathbf{R} = \text{ENC}(\mathcal{A}, X; \Theta_{Enc})$$

- **Knowledge Graph decoder** $\text{DEC}_{\Theta_{Dec}}$:

$$\text{DEC}_{\Theta_{Dec}} : \mathbb{R}^{d_e} \times \mathbb{R}^{d_r} \times \mathbb{R}^{d_e} \rightarrow \mathbb{R}$$

The decoder is parameterized by $\Theta_{Dec}$ and, given the embeddings of the head entity $\mathbf{h} \in \mathbb{R}^{d_e}$, the relation $\mathbf{r} \in \mathbb{R}^{d_r}$, and the tail entity $\mathbf{t} \in \mathbb{R}^{d_e}$, scores the corresponding triple $(h, r, t)$ to represent the likelihood or strength of the interaction.

Formally:

$$s(h, r, t) = \text{DEC}(\mathbf{h}, \mathbf{r}, \mathbf{t}; \Theta_{Dec})$$

The scoring function $s(h, r, t)$ can be instantiated in different ways depending on the type of model.

The encoder and decoder are trained using a loss function $\mathcal{L}$ that compares the scores of positive and negative triples.

Positive triples correspond to observed triples in the KG, while negative triples represent invalid triples that are not present in the KG. The model are trained to distinguish between positive and negative triples, assigning higher scores to valid triples and lower scores to invalid ones.

Formally, the loss function can be defined as:

$$\mathcal{L} = \sum_{(h,r,t)\in E^+} \sum_{(h',r,t')\in E^-} \ell(s(h, r, t), s(h', r, t'))$$

where:

- $E^+$ is the set of positive triples,
- $E^-$ is the set of negative triples,
- $\ell(\cdot, \cdot)$ is a cost function that compares the scores of the positive and negative triples.

##### *S1.3.1. Shallow and Deep Embeddings*

The encoder can take the form of a simple embedding look-up table, associating each entity and relation with a pre-defined vector representation. In this case, the encoder can be defined as:

$$\mathbf{Z}, \mathbf{R} = \text{ENC}(\Theta_{Enc})$$

This approach results in *shallow embeddings*, where the encoder does not incorporate the graph structure. Instead, the topology of the knowledge graph is only accounted for at the decoder stage.

In contrast, other models employ encoders that integrate the graph structure during the encoding process by utilizing $\mathcal{A}$, and optionally the entities feature matrix $X$. These encoders capture the structural relationships between entities, often relying on Graph Neural Networks (GNNs) models. This results in *deep embeddings*, where the encoder generates representations that reflect the structural roles of entities within the graph. GNN encoders can be used in combination with shallow embedding decoders to perform link prediction in KGs.

#### S1.4. Encoder used in this study

**GAT** (2) is a graph neural network encoder based on an attention mechanism. Each entity embedding is updated by aggregating the embeddings of its neighbors within each relation type, weighted by learned attention coefficients that quantify each neighbor's relevance. The resulting per-relation representations are then summed across relation types. The GAT encoder can be coupled and trained with any of the decoders described below.

**GCN** (3) is a graph neural network encoder that, like GAT, aggregates information at two levels but uses a fixed, non-learned operator for the intra-relation aggregation. Each entity embedding is updated by aggregating the embeddings of its neighbors within each relation type using a fixed pooling operator, for which we evaluated two schemes, mean and max; the resulting per-relation representations are then summed across relation types. The GCN encoder can be coupled and trained with any of the decoders described below.

#### S1.5. Decoders used in this study

KGE decoders can be divided into two main categories: translational and bilinear models (4). Translational models represent relations as transformations in the embedding space, typically aiming to capture the relationship between entities through geometric operations like translations or rotations in the embedding space. In contrast, bilinear models use multiplicative interactions between embeddings, often through tensor operations, to capture complex relational patterns.

##### *S1.5.1. Translational Models*

**TransE** (5) is the simplest translational model. It represents relations as translation vectors and assumes that $\mathbf{h} + \mathbf{r} \approx \mathbf{t}$ for valid triples. The scoring function is defined as the negative distance between the translated head and the tail:

$$s(h, r, t) = -\|\mathbf{h} + \mathbf{r} - \mathbf{t}\|_p$$

where $\|\cdot\|_p$ denotes the $L_1$ or $L_2$ norm.

**TransD** (6) extends TransE by introducing entity- and relation-specific projection vectors that dynamically map entities into a relation-specific space before applying the translation. This allows the model to better handle complex relational patterns and to support distinct embedding dimensions for entities ($d_e$) and relations ($d_r$).

##### *S1.5.2. Bilinear Models*

**RESCAL** (7) represents each relation as a full matrix $\mathbf{M}_r \in \mathbb{R}^{d_e \times d_e}$ and scores a triple through a bilinear product:

$$s(h, r, t) = \mathbf{h}^\top \mathbf{M}_r \mathbf{t}$$

**DistMult** (8) simplifies RESCAL by restricting relation matrices to be diagonal and thus constraining the model to symmetric relations only.

**ComplEx** (9) extends DistMult to the complex domain, embedding entities and relations as vectors in $\mathbb{C}^d$. It can model both symmetric and antisymmetric relations while retaining the parameter efficiency of DistMult.

**HolE** (10) composes head and tail embeddings through circular correlation and scores the triple by the dot product between the resulting vector and the relation embedding.

**ANALOGY** (11) imposes analogical structure on relation matrices by constraining them to be normal and mutually commutative. It is a generalization of DistMult, ComplEx, and HolE within a single framework.

#### S1.6. Negative Sampling

While positive triples can be directly extracted from the KG, selecting negative triples is more challenging, as most KGs only contain true facts. A common approach to generate negative triples is Negative Sampling, where invalid triples are created by corrupting positive ones. This is done by randomly replacing the head (or tail) entity in a triple while keeping the relation and the other entity fixed.

It has been shown that the quality of generated negative triples has a direct impact on the performance of KGE models in a myriad of downstream tasks (12), including link prediction.

#### S1.7. Evaluation Metrics

Link prediction is evaluated through a ranking procedure: each evaluated triple $(h, r, t)$ is used to predict the tail given $(h, r)$ and the head given $(r, t)$. For each prediction, candidate entities are ranked from most to least plausible according to the model's scoring function, and the rank of the true entity is recorded. We report two standard metrics over these ranks:

**Mean Reciprocal Rank (MRR).** The average of the reciprocal ranks of the true entity:

$$\text{MRR} = \frac{1}{N} \sum_{i=1}^{N} \frac{1}{\text{rank}_i},$$

where $N$ is the total number of predictions and $\text{rank}_i$ is the rank of the true entity for prediction $i$.

**Hits@$k$.** The proportion of predictions for which the true entity is ranked within the top $k$ candidates.

Both metrics are bounded between 0 and 1, with higher values indicating better performance.

Following the standard *filtered* setting, candidates ranked above the true entity that correspond to other valid triples in the KG are removed from the ranking, to avoid penalizing the model for making a correct prediction.

##### S1.8. Semantically redundant relations

Let $T_r$ denote the set of head-tail pairs in relation $r$, defined as:

$$T_r = \{(h,t) \mid (h,r,t) \in \mathcal{G}\}.$$

Let $T_r^{-1}$ denote the set of tail-head pairs in relation $r$, defined as:

$$T_r^{-1} = \{(t,h) \mid (h,r,t) \in \mathcal{G}\}.$$

Let $|r|$ represent the number of instance triples in relation $r$.

Two relations $r_1$ and $r_2$ are considered **near-duplicate relations** if they satisfy the following condition:

$$\frac{|T_{r_1} \cap T_{r_2}|}{|r_1|} > \theta_1 \quad \text{and} \quad \frac{|T_{r_1} \cap T_{r_2}|}{|r_2|} > \theta_2, \tag{S1}$$

Similarly, two relations $r_1$ and $r_2$ are considered **(near)-reverse-duplicate relations** if they satisfy the following condition:

$$\frac{|T_{r_1} \cap T_{r_2}^{-1}|}{|r_1|} > \theta_1 \quad \text{and} \quad \frac{|T_{r_1} \cap T_{r_2}^{-1}|}{|r_2|} > \theta_2, \tag{S2}$$

##### S1.9. Cartesian product relations

A relation $r$ is considered a **Cartesian product relation** if it satisfies the following condition:

$$\frac{|T_r|}{|S_r| \cdot |O_r|} > \theta \tag{S3}$$

where $S_r$ and $O_r$ represent the sets of unique heads and tails for the relation $r$, respectively, formally defined as:

$$S_r = \{h \mid \exists (h,r,t) \in \mathcal{G}\}, \quad O_r = \{t \mid \exists (h,r,t) \in \mathcal{G}\}.$$

#### S2. Supplementary Results

##### S2.1. Sensitivity Analysis on $\theta_1$, $\theta_2$ and $\theta$ Thresholds

To detect semantically redundant relations in our KGs, we followed Akrami et al. (13) and used a threshold of 0.8 on head-tail pair overlap (for near-duplicate and reverse-duplicate relations) and on head-tail Cartesian product density (for Cartesian product relations). To assess the sensitivity of the results to this threshold, we ran the DL1 detection and control procedure on all three KGs (ShepherdKG, BioKG, HetionetKG) using both the original threshold of 0.8 and a threshold of 0.5, which can potentially increase the number of relations considered as semantically redundant.

No semantically redundant (near-duplicate, reverse-duplicate, or Cartesian product relations) were detected on ShepherdKG or Hetionet at either threshold. On BioKG, the DPI and DRUG_TARGET relations, both connecting drug nodes to protein nodes, were flagged as near-duplicate relations at 0.5, but no relation was flagged at 0.8; no reverse-duplicate or Cartesian product relations were detected at either threshold.

##### S2.2. DL1 detection and control procedure on BioKG and HetionetKG

We applied the DL1 detection and control procedure described in Materials and Methods for ShepherdKG to BioKG and HetionetKG. Below, we describe the KG-specific outcomes of each step.

**BioKG** comprises $105,522$ entities and 19 relation types. Step 1 did not remove any triples, as no relation in BioKG was the reverse or a duplicate of another. At Step 2, using $\theta_1$, $\theta_2$, and $\theta$ set to 0.8, we did not find any redundant relationship types, but using $\theta_1$, $\theta_2$, and $\theta$ set to 0.5, we identified two near-duplicate relations: *DPI* (drug-protein interaction) and *DRUG_TARGET*, which were added to the list of known redundant relations. We continued the analysis with $\theta_1$, $\theta_2$, and $\theta$ set to 0.5. At Step 3, two undirected relationship types were made directed: drug-drug interactions (*DDI*) and protein-protein interactions (*PPI*). For these relations, the reverse was included with a different relation name (e.g., *DDI_rev*) to avoid introducing symmetric relationships.

**HetionetKG** comprises $45,158$ entities and 24 relation types. At Step 1, $1,689$ duplicated triples were removed from the gene-regulates-gene relation type (*GrG*). At Step 2, using $\theta_1$, $\theta_2$, and $\theta$ set to 0.8 or 0.5, no semantically redundant or Cartesian product relations were identified. At Step 3, four undirected relation types were made directed: disease-resembles-disease (*DrD*), compound-resembles-compound (*CrC*), gene-interacts-gene (*GiG*), and gene-covaries-gene (*GcG*). For these relations, the reverse was included with a different relation name to avoid introducing symmetric relationships.

For both KGs, the reverse triples introduced at Step 3 created a semantic redundancy that was removed at Step 4 of the DL1 detection and control procedure for the KG controlling for data leakage, but retained for the KG where data leakage was not controlled.

For BioKG, Step 4 also removed $4,928$ triples from the training set due to the semantic redundancy of the *DPI* and *DRUG_TARGET* relations.

#### S2.3. DL1 leakage operates through different channels in decoder-only and GNN models

##### *S2.3.1. How DL1 leakage works in decoder-only models*

DL1 stems from redundancy between relation types. Let us consider two redundant relations $r$ and $r'$ that are near-duplicates or near-reverse-duplicates. DL1 can affect a decoder-only model through two learning mechanisms that act jointly to produce the performance inflation observed in our decoder-only models.

First, on the relation side, what we call the *relation equivalence*: from the many redundant triples seen in training (e.g. $(a, r, b)$ and $(a, r', b)$), the model learns to give $r$ and $r'$ linked representations, so that the decoder scores a triple and its redundant counterpart highly.

Second, on the entity side, what we call *per-pair memorization*: every triple seen in training also pulls the embeddings of its head and tail toward a configuration in which they score highly under the relation of that triple.

The relation equivalence mechanism is in fact an expected behavior: $r$ and $r'$ encode the same underlying fact, and a model is expected to represent equivalent facts consistently. Therefore, the problem is not that the model learns this equivalence, but that the training/validation/test split can turn it into a shortcut using per-pair memorization if not handled carefully (i.e. if DL1 is not controlled). Indeed, DL1 occurs only when the split separates a redundant pair. Suppose $(a, r, b)$ is held out for evaluation in the test set, while its redundant counterpart $(a, r', b)$ (or $(b, r', a)$ in the case of near-reverse-duplicates) remains in training. The two mechanisms then act together: per-pair memorization drives the embeddings of $a$ and $b$ into the high-scoring configuration under $r'$, and because $r$ and $r'$ have been given linked representations, that same configuration also scores $(a, r, b)$ highly. The held-out evaluation triple is thus recoverable jointly from the entity embeddings shaped by its redundant counterpart and from the learned equivalence between relations $r$ and $r'$.

Taken together, we call these two mechanisms the *decoder DL1 channel*. The without-DL1 setting does not remove the redundancy globally. It removes, from the training set, only the redundant counterpart of triples that fall in validation or test. The model still learns the equivalence between $r$ and $r'$ from all the redundant pairs that legitimately coexist in training. However, for an evaluation triple, the model no longer receives the redundant counterpart that would shape $a$ and $b$ into the high-scoring configuration.

##### *S2.3.2. Two DL1 leakage channels in GNN models*

A GNN encoder builds the embedding of a node in two stages: it first aggregates, within each relation type, the embeddings of neighbors connected through that relation (intra-relation aggregation), and then combines these relation-specific representations across relation types (inter-relation aggregation), typically using fixed operators such as mean, sum, max, or using learned attention in the case of GAT.

Since a GNN encoder is combined with a decoder, the decoder DL1 channel described above still applies: the model still scores the redundant training triples, so the relation equivalence is still learned and can still be exploited on an evaluation pair whose counterpart was left in training. However, the entity embeddings are no longer fully-free parameters but are produced by aggregation, so per-pair memorization can no longer place $a$ and $b$ as freely as in a decoder-only model. We therefore expect the decoder DL1 channel to survive in GNN models, but in attenuated form.

In addition, GNN models introduce a second, new potential DL1 channel. A redundant counterpart such as $(b, r', a)$ is not only scored by the decoder; it is also an edge of the message-passing graph, so it places $b$ in the neighborhood of $a$ and lets $b$ contribute to the embedding of $a$ during encoding. We call this the *message-passing DL1 channel*. It has no equivalent in decoder-only models, where embeddings are fully-free parameters and no aggregation occurs. The question is whether this contribution can create similar performance inflation as the decoder DL1 channel. The key difference lies in the learning signal. The decoder DL1 channel relies on pair-specific supervision: the training triple $(b, r', a)$ explicitly teaches the model that this particular pair should receive a high score. The resulting gradients directly adjust the representations involved in that pair, enabling the decoder to memorize it. Message passing provides no such supervision. During encoding, $b$ contributes to the representation of $a$ only as one of its neighbors connected by relation $r'$. The encoder is never informed that $b$ is the redundant counterpart of a held-out evaluation triple, as that information exists only in the test set. Consequently, while an aggregation mechanism may assign different weights to different neighbors (for example through attention), nothing in the encoder encourages it to treat $b$ differently. As a result, the information carried by $b$ may indeed influence the representation of $a$, but this influence remains a neighborhood-level structural signal rather than the pair-specific memorization responsible for DL1 in decoder-only models. Aggregation can also attenuates this structural signal by mixing information from multiple neighbors within each relation and subsequently across relation types, diluting the contribution of any single neighbor. Whether this diffuse structural contribution meaningfully inflates performance is not guaranteed: lacking pair-specific supervision, the message-passing cannot recover the held-out pair as directly as the decoder DL1 channel.

Among the GNN models evaluated, GCNmean displays a counter-intuitive pattern: its performance is systematically lower in the with-DL1 setting than in the without-DL1 setting, in contrast to all other models for which DL1 leakage inflates scores. This pattern can be explained by the interaction between mean aggregation and graph density. Mean aggregation computes a node's representation as the arithmetic mean of its neighbors' embeddings within each relation type. As a consequence, each individual neighbor contributes a fraction $1/k$ of the aggregated signal, where $k$ is the neighborhood size. On biomedical KGs, $k$ is often large, diluting the contribution of any single neighbor. DL1 leakage further increases graph density by retaining redundant counterpart triples in the training set, which enlarges node neighborhoods and amplifies this dilution effect. The useful structural signal carried by any individual neighbor is therefore more severely attenuated in the with-DL1 graph than in the without-DL1 graph, explaining the observed performance inversion. Max aggregation, by contrast, does not average over the neighborhood, so strong signals are preserved rather than diluted, which may make GCNmax robust to graph densification, consistent with its outperforming GCNmean in all experimental conditions.

#### S2.4. Degree can be encoded in entity embeddings without driving link prediction performance

##### *S2.4.1. Degree can be encoded in entity embeddings but does not drive prediction in decoder-only models*

DL2 stems from degree imbalance within a single relation type $r$: a small number of hub entities concentrate most of the connections, appearing as the tail of $r$ with many distinct heads. The concern is that a model might rank these hubs highly not because they are the correct answer to a given query, but simply because they are frequent, using degree as an illegitimate feature.

We examine whether a decoder-only model can exploit degree, in two steps: whether degree is encoded in the embeddings at all, and, if so, whether that encoding can translate into a link-prediction advantage.

A natural place for degree to be encoded is the embedding norm. Every training triple pulls the embeddings of its head and tail toward a configuration that scores highly under its relation. An entity that is the tail of $r$ with many heads accumulates many such updates, and, without any constraint, its norm grows with its degree. Whether this happens depends on how each model treats the entity norm during training, which splits our decoder-only models into two groups. ComplEx and ANALOGY leave the entity embedding unconstrained, so their norms are free to grow. DistMult, RESCAL, HolE, TransE, and TransD instead renormalize entity embeddings after each update, leaving every entity embedding with norm = 1. Thus, for such embedding models, degree is, by construction, not encoded in the entity embedding through the norm. So degree may or may not be written into the norm, depending on the model. To evaluate that, we measure the correlation between each entity's embedding $L_2$ norm and (i) its global degree (over all relations, in and out degree); (ii) its drug degree within the indication relation (the number of diseases a drug is indicated for); and (iii) its disease degree within the indication relation (the number of drugs indicated for the disease).

Another place where degree can be encoded is in the direction of the entity embeddings. To isolate direction, we project every embedding onto the unit sphere by dividing it by its $L_2$ norm. Direction is a vector, whereas correlating it with degree requires a scalar per entity. We obtain this scalar by measuring how closely an entity's direction aligns with a reference direction, computed as the cosine to the centroid (mean direction) of a reference set of entities. This reference set depends on the degree being correlated: (i) for global degree, the reference is the mean direction of all entities; (ii) for drug degree within the indication relation, the reference is the mean direction of the drugs; and (iii) for disease degree within the indication relation, the reference is the mean direction of the diseases. A positive correlation then means that higher-degree entities point more toward their reference direction. A negative correlation means that higher-degree entities point away from the reference. A correlation close to zero means orientation carries no degree information.

We performed the norm and direction analyses on the original and permuted versions of ShepherdKG and BioKG (Supplementary Figures 18 and 19). We show that global degree is broadly encoded: in the norm for the unconstrained models (ANALOGY and ComplEx) and, by construction, not for the others; and in the direction for all decoder-only models, though to a lesser extent for TransD.

The degree relevant to DL2, however, is not the global degree but the entity's degree within the permuted relation (i.e. the indication relation). On ShepherdKG, drug degree within the indication relation is not encoded in the norm for any decoder-only model, whereas disease degree carries a moderate norm signal for the two unconstrained models (ANALOGY and ComplEx, with a disease-degree-to-norm correlation ranging from 0.39 to 0.46). In the direction, drug degree is likewise not encoded in decoder-only models, while the correlation between direction and disease degree varies from model to model and between the original and permuted KGs. Thus, on ShepherdKG, whether or not a given decoder-only model encodes indication degree, they predict the true indications relatively well and fail on the permuted graph (Main Text Figure 2). For several models the disease-degree-to-direction correlation is even higher on the permuted ShepherdKG, i.e. degree is better encoded in the permuted ShepherdKG. These findings show that, even though decoder-only models can encode degree in their embeddings, this encoding explains neither their accuracy on the original ShepherdKG nor the performance gap between the original and permuted graphs.

On BioKG, degree encoding is stronger. The unconstrained decoder-only models (ANALOGY and ComplEx) encode indication degree substantially in the norm, on both the drug side (correlation $\geq 0.46$) and, more strongly, the disease side (correlation $\geq 0.73$). In the direction, as on ShepherdKG, the relation-specific correlations vary across models and graph versions. Here global and indication degree are clearly encoded, yet decoder-only models reach only poor indication-prediction performance on both the original and permuted BioKG (Supplementary Figure 9). Strong degree encoding is thus not enough to yield good predictions, as decoder-only models fail to predict indication relations in BioKG.

Taken together, the two graphs dissociate degree encoding from predictive performance in opposite directions: on ShepherdKG, decoder-only models predict indications relatively well while encoding indication degree only weakly and inconsistently; on BioKG, they encode it strongly yet predict poorly. Encoding degree is therefore neither necessary nor sufficient to explain the performance of decoder-only models for indication relations.

##### *S2.4.2. Degree-correlated features may drive prediction in GNN-based models*

Decoder-only and GNN-based models are trained from exactly the same triples, and both can in principle learn structural properties present in the graph. The difference lies in the representation mechanism. In decoder-only models, structural properties can only emerge indirectly, through the optimization of independent entity embeddings under the decoder objective. In GNN-based models, they are computed explicitly by message passing, making neighbourhood structure an intrinsic component of every node representation.

BioKG illustrates this distinction. For BioKG, decoder-only models encode degree, yet fail to predict original or permuted indications. GNN encoders encode degree as well (Supplementary Figure 19), yet, unlike decoder-only models, they retain strong indication-prediction performance even on the permuted KG (Supplementary Figure 9). This indicates that they may exploit structural information that survives the permutation.

Our DL2 construction preserves the indication-relation degree distribution by design, making degree one such surviving signal. Degree alone, however, may not account for the retained performance of GNN-based models on the permuted BioKG: decoder-only models also encode degree, yet show low performance in predicting (original or permuted) indications on BioKG. Moreover, because message passing also exposes other structural properties correlated with degree, our experiments do not allow us to attribute the retained performance of GNN-based models to degree alone. Rather, they show that, on BioKG, GNN encoders may exploit other structural features preserved by the permutation, whereas decoder-only models do not.

### S3. Supplementary Tables

**Supplementary Table S1.** Entity and Relationship Counts in the ShepherdKG Knowledge Graph

**(a) Entity types**

| Type | Count |
|---|---|
| biological_process | 28642 |
| gene/protein | 27610 |
| disease | 21258 |
| effect/phenotype | 15874 |
| anatomy | 14033 |
| molecular_function | 11169 |
| drug | 7946 |
| cellular_component | 4176 |
| pathway | 2516 |
| exposure | 802 |
| Total | 134026 |

**(b) Relationship types**

| Relation Type | Count |
|---|---|
| drug_drug | 2672628 |
| protein_present_anatomy | 1518203 |
| protein_protein | 321075 |
| disease_phenotype_positive | 204766 |
| protein_bioprocess | 144805 |
| disease_protein | 86299 |
| protein_cellcomp | 83402 |
| drug_effect | 79137 |
| protein_molfunc | 69530 |
| bioprocess_bioprocess | 52886 |
| protein_pathway | 42646 |
| disease_disease | 35171 |
| contraindication | 28884 |
| drug_protein | 25653 |
| phenotype_phenotype | 21925 |
| protein_absent_anatomy | 19887 |
| anatomy_anatomy | 14032 |
| molfunc_molfunc | 13574 |
| phenotype_protein | 10518 |
| indication | 8533 |
| cellcomp_cellcomp | 4845 |
| pathway_pathway | 2535 |
| off-label use | 2457 |
| exposure_exposure | 2140 |
| exposure_disease | 1795 |
| exposure_bioprocess | 1625 |
| disease_phenotype_negative | 1483 |
| exposure_protein | 1212 |
| exposure_molfunc | 45 |
| exposure_cellcomp | 10 |
| Total | 5471699 |

**Supplementary Table S2.** Entity and Relationship Counts in the BioKG Knowledge Graph

**(a) Entity types**

| Type | Count |
|---|---|
| protein | 59204 |
| pathway | 14779 |
| complex | 13246 |
| drug | 8805 |
| disease | 5812 |
| genetic_disorder | 3678 |
| Total | 105524 |

**(b) Relationship types**

| Relation Type | Count |
|---|---|
| DDI | 1334085 |
| PROTEIN_PATHWAY_ASSOCIATION | 250118 |
| PPI | 113817 |
| PROTEIN_DISEASE_ASSOCIATION | 109276 |
| MEMBER_OF_COMPLEX | 87452 |
| DRUG_DISEASE_ASSOCIATION | 66867 |
| DPI | 28033 |
| COMPLEX_IN_PATHWAY | 21125 |
| COMPLEX_TOP_LEVEL_PATHWAY | 15639 |
| DRUG_TARGET | 15231 |
| DRUG_PATHWAY_ASSOCIATION | 5114 |
| DRUG_ENZYME | 5077 |
| DISEASE_GENETIC_DISORDER | 4882 |
| RELATED_GENETIC_DISORDER | 3894 |
| DISEASE_PATHWAY_ASSOCIATION | 3544 |
| DRUG_TRANSPORTER | 3040 |
| DRUG_CARRIER | 804 |
| Total | 2067998 |

**Supplementary Table S3.** Entity and Relationship Counts in the HetionetKG Knowledge Graph

**(a) Entity types**

| Type | Count |
|---|---|
| gene | 19145 |
| biological process | 11381 |
| side effect | 5701 |
| molecular function | 2884 |
| pathway | 1822 |
| compound | 1538 |
| cellular component | 1391 |
| symptom | 415 |
| anatomy | 400 |
| pharmacologic class | 345 |
| disease | 136 |
| Total | 45158 |

**(b) Relationship types**

| Relation Type | Count |
|---|---|
| gene_participates_biological_process | 559504 |
| anatomy_expresses_gene | 526407 |
| gene_regulates_gene | 265672 |
| gene_interacts_gene | 147164 |
| compound_causes_side_effect | 138944 |
| anatomy_downregulates_gene | 102240 |
| anatomy_upregulates_gene | 97848 |
| gene_participates_molecular_function | 97222 |
| gene_participates_pathway | 84372 |
| gene_participates_cellular_component | 73566 |
| gene_covaries_gene | 61690 |
| compound_downregulates_gene | 21102 |
| compound_upregulates_gene | 18756 |
| disease_associates_gene | 12623 |
| compound_binds_gene | 11571 |
| disease_upregulates_gene | 7731 |
| disease_downregulates_gene | 7623 |
| compound_resembles_compound | 6486 |
| disease_localizes_anatomy | 3602 |
| disease_presents_symptom | 3357 |
| pharmacologic_class_includes_compound | 1029 |
| compound_treats_disease | 755 |
| disease_resembles_disease | 543 |
| compound_palliates_disease | 390 |
| Total | 2250197 |

#### S4. Supplementary Figures

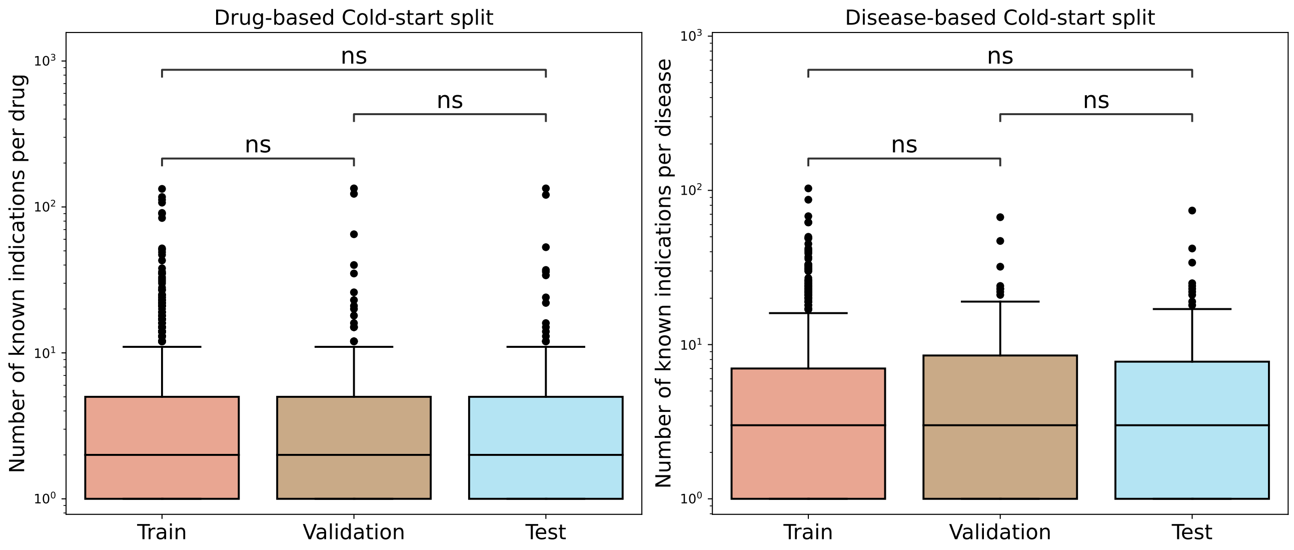


**Supplementary Figure S1.** Distribution, in **ShepherdKG**, of the number of known indications per drug (left) and per disease (right) across the train, validation, and test sets in drug-based and disease-based cold-start settings, respectively. Statistical significance was assessed using the Mann-Whitney-Wilcoxon test (two-sided). Annotations indicate significance levels: $ns$: $5.00 \times 10^{-2} < p \leq 1$.

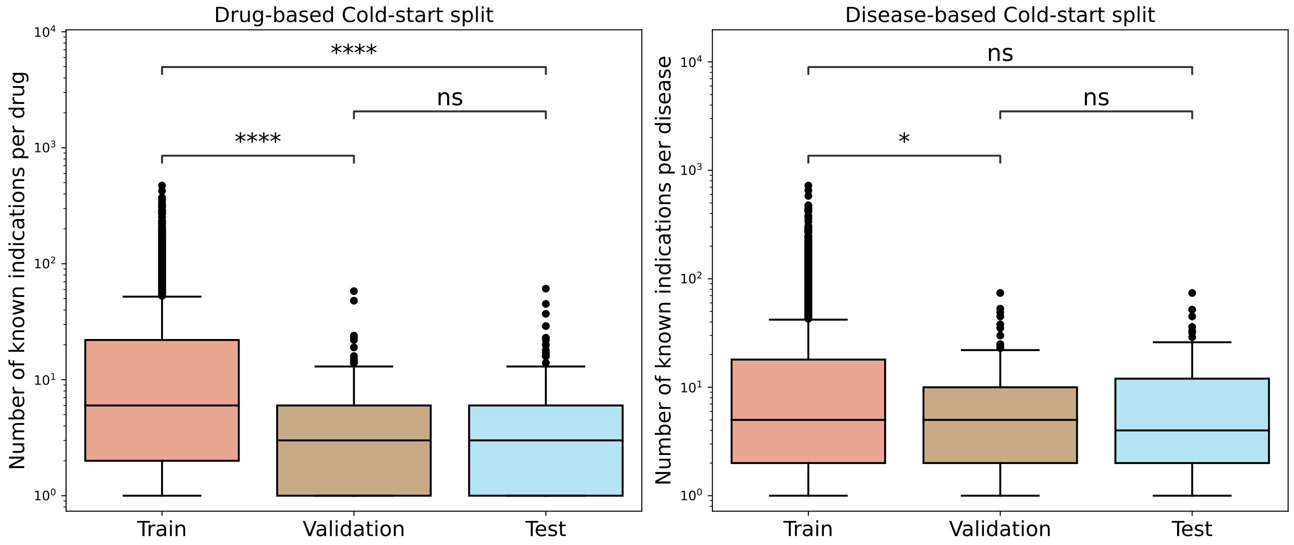


**Supplementary Figure S2.** Distribution, in **BioKG**, of the number of known indications per drug (left) and per disease (right) across the train, validation, and test sets in drug-based and disease-based cold-start settings, respectively. Note that, in contrast to ShepherdKG and HetionetKG (Supplementary Figures S1 and S3), the validation and test sets have slightly lower indication frequencies than the training set. This stems from the structure of BioKG, where many drugs and diseases appear exclusively in the indication (DRUG_DISEASE_ASSOCIATION) relation. Since no other training triplets exist for these entities, the model cannot learn any embedding for them outside of the training set; they are therefore kept in training only. This reduces the pool of entities eligible for cold-start splits, introducing a slight bias toward less-indicated entities in the validation and test sets. Statistical significance was assessed using the Mann-Whitney-Wilcoxon test (two-sided). Annotations indicate significance levels: $ns$: $5.00 \times 10^{-2} < p \leq 1$. $*$: $1.00 \times 10^{-2} < p \leq 5.00 \times 10^{-2}$; $****$: $p \leq 1.00 \times 10^{-4}$.

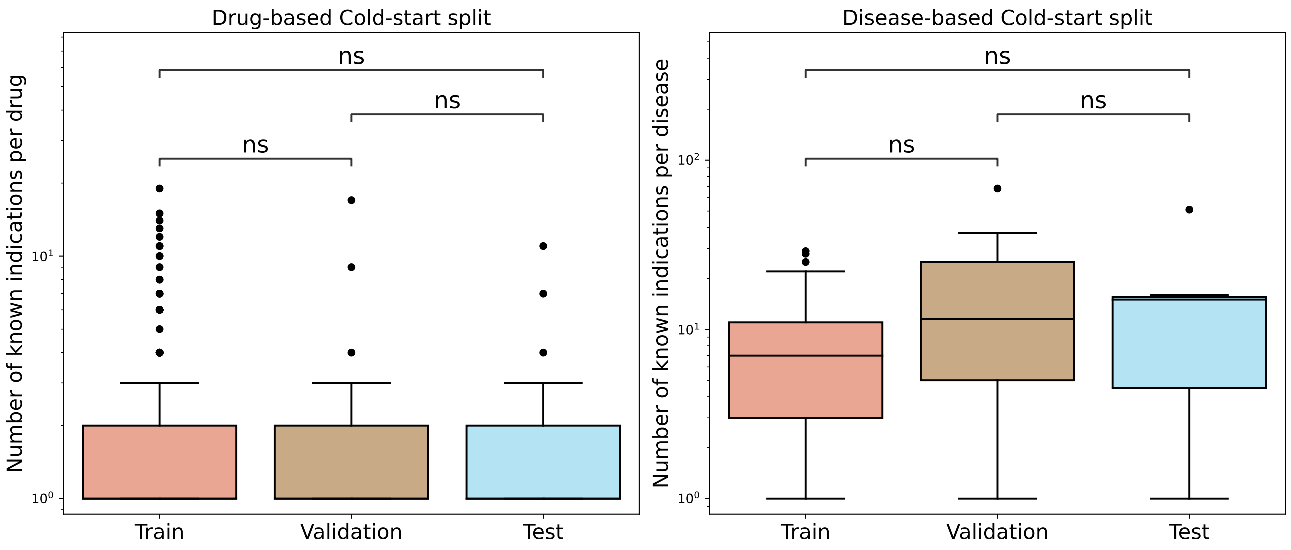


**Supplementary Figure S3.** Distribution, in **HetionetKG**, of the number of known indications per drug (left) and per disease (right) across the train, validation, and test sets in drug-based and disease-based cold-start settings, respectively. Statistical significance was assessed using the Mann-Whitney-Wilcoxon test (two-sided). Annotations indicate significance levels: $ns$: $5.00 \times 10^{-2} < p \leq 1$.

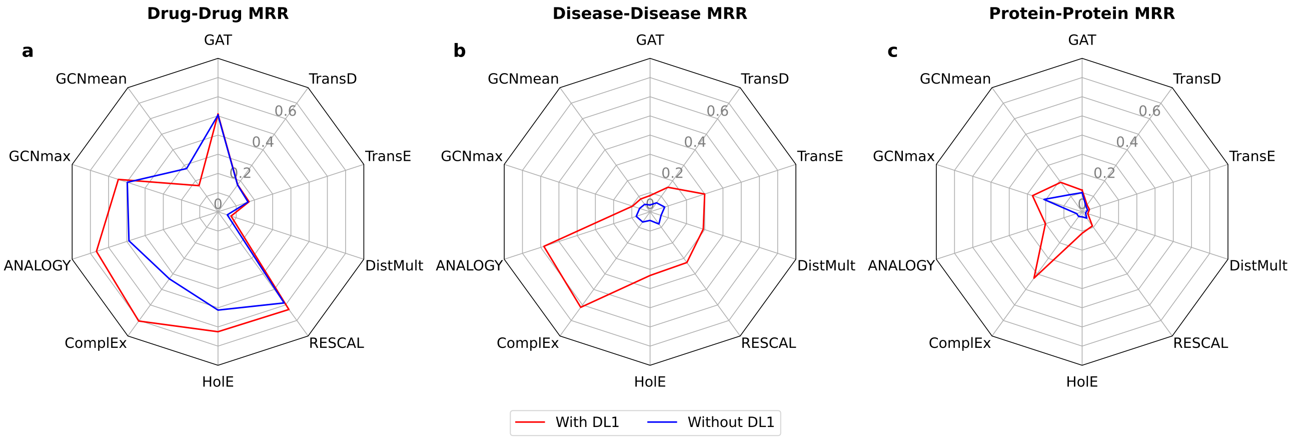


**Supplementary Figure S4.** Impact of Training-Test Set Separation on KGE Model Performance per Redundant Relation Type (DL1) on **ShepherdKG**. **(a)** Drug-Drug MRR: model performance on drug-drug interaction triples (average of MRR on forward and inverse relations). **(b)** Disease-Disease MRR: model performance on disease-disease triples (average of MRR on forward and inverse relations). **(c)** Protein-Protein MRR: model performance on protein-protein interaction triples (average of MMR on forward and inverse relations). Performance shown with (red) and without (blue) DL1.

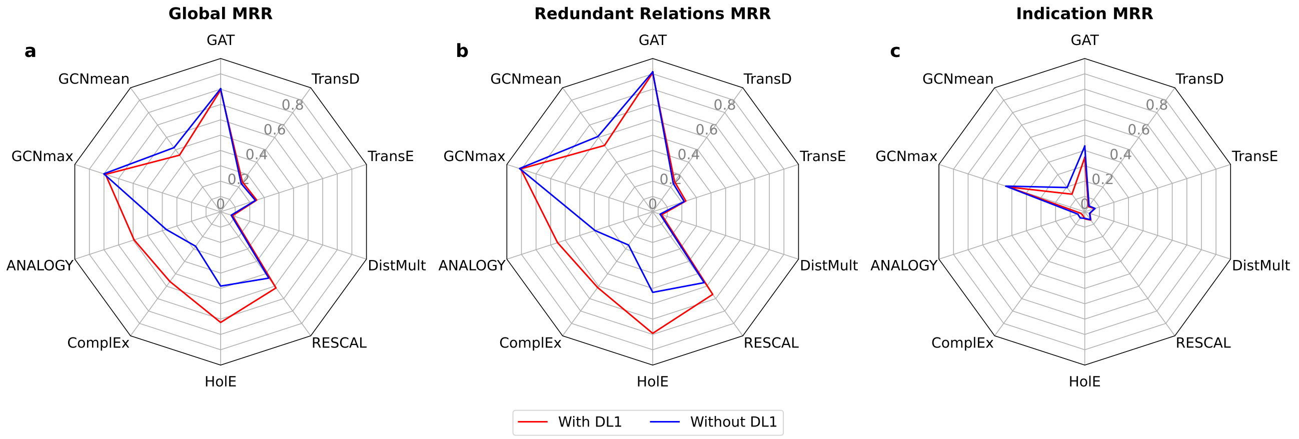


**Supplementary Figure S5.** Impact of Training-Test Set Separation on KGE Model Performance (DL1) on **BioKG**. **(a)** Global MRR: performance of the models across all relations in the test set. **(b)** MRR for Redundant Relations: Model performance on the relations flagged as redundant by the DL1 detection and control procedure. **(c)** MRR for the Indication Relation: Model performance on indication triples (DRUG_DISEASE_ASSOCIATION). Performance shown with (red) and without (blue) DL1.

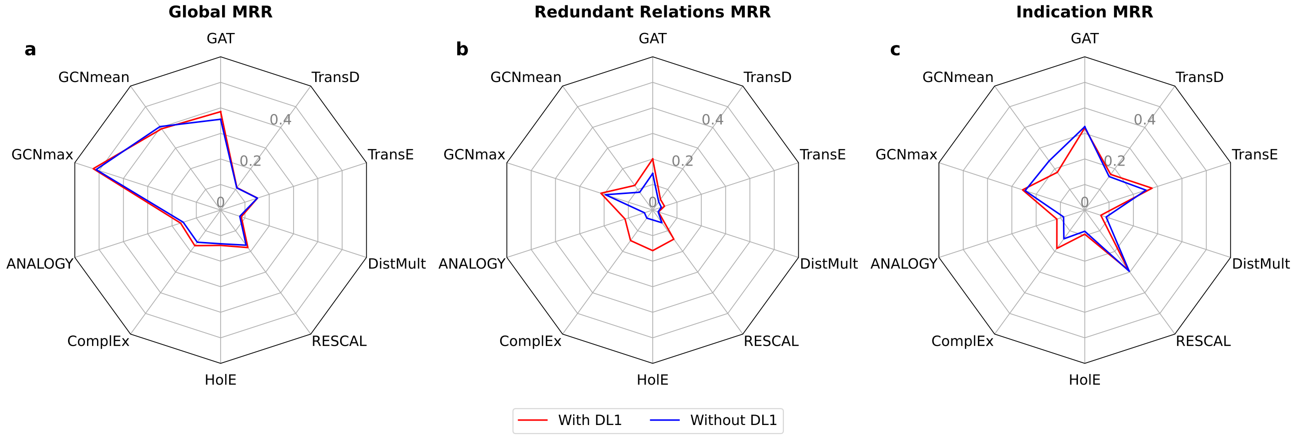


**Supplementary Figure S6.** Impact of Training-Test Set Separation on KGE Model Performance (DL1) on **HetionetKG**. **(a)** Global MRR: performance of the models across all relations in the test set. **(b)** MRR for Redundant Relations: Model performance on the relations flagged as redundant by the DL1 detection and control procedure. **(c)** MRR for the Indication Relation: Model performance on indication triples (compound_treats_disease). Performance shown with (red) and without (blue) DL1.

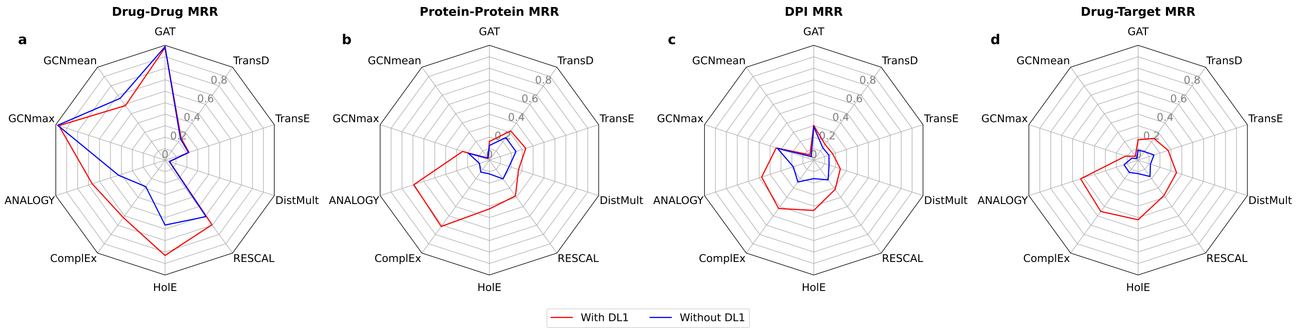


**Supplementary Figure S7.** Impact of Training-Test Set Separation on KGE Model Performance per Redundant Relation Type (DL1) on **BioKG**. **(a)** Drug-Drug MRR: model performance on drug-drug association triples (average of MRR on forward and inverse relations). **(b)** Protein-Protein MRR: model performance on protein-protein interaction triples (average of MRR on forward and inverse relations). **(c)** DPI MRR: model performance on drug-protein interaction triples. This relation is semantically redundant with the Drug-Target relation. **(c)** Drug-Target MRR: model performance on drug-target protein interaction triples. This relation is semantically redundant with the DPI relation. Performance shown with (red) and without (blue) DL1.

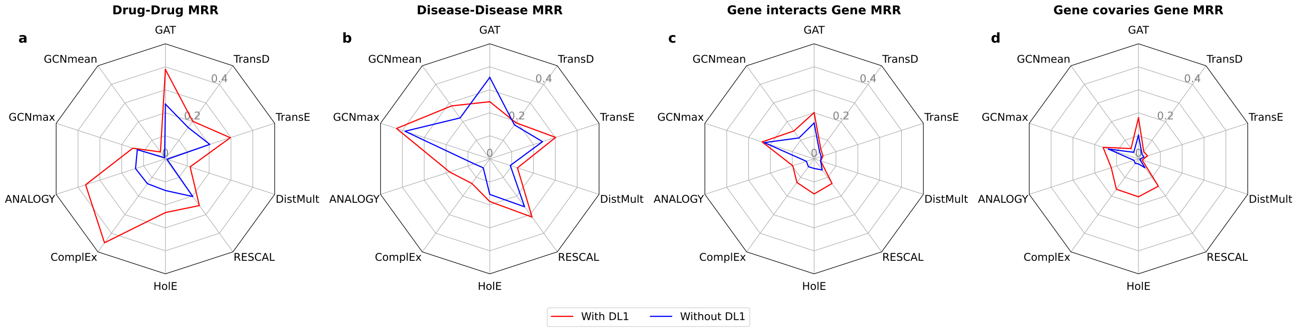


**Supplementary Figure S8.** Impact of Training-Test Set Separation on KGE Model Performance per Redundant Relation Type (DL1) on **HetionetKG**. **(a)** Drug-Drug MRR: model performance on drug-drug association triples (average of MRR on forward and inverse relations). **(b)** Disease-Disease MRR: model performance on disease-disease triples (average of MRR on forward and inverse relations). **(c)** Gene-Gene interaction MRR: model performance on gene-interacts-gene triples (average of MRR on forward and inverse relations). **(d)** Gene-Gene covariation MRR: model performance on gene-covaries-gene triples (average of MRR on forward and inverse relations). Performance is shown with (red) and without (blue) DL1.

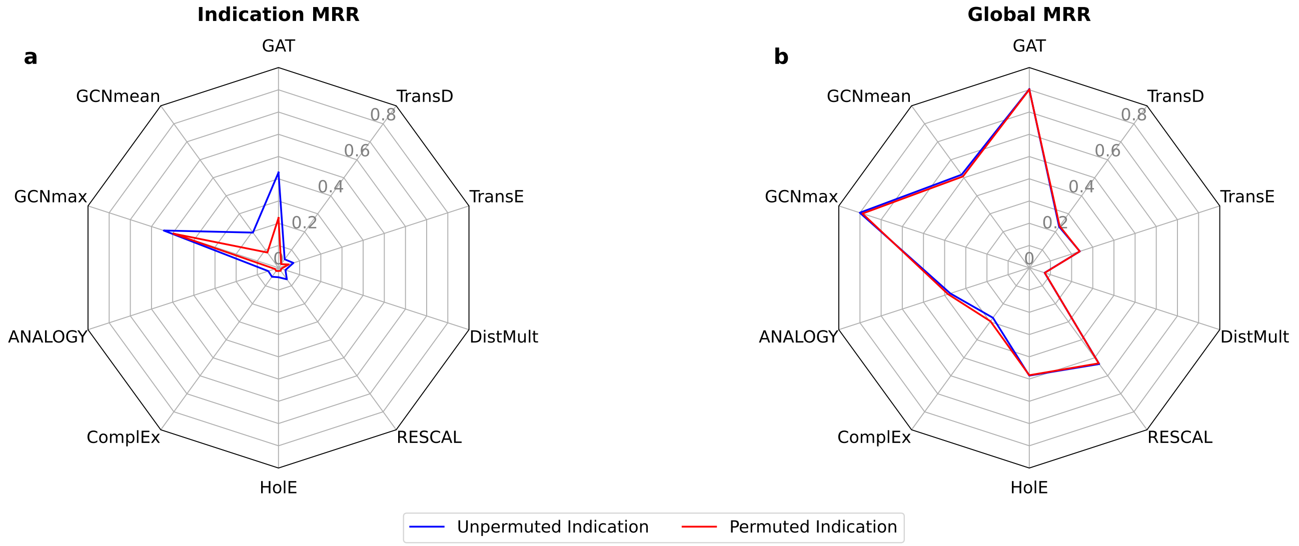


**Supplementary Figure S9.** Assessment of node degree exploitation as an illegitimate feature (DL2) in indication prediction on **BioKG**. **(a)** MRR for the Indication Relation: Model performance on indication (DRUG_DISEASE_ASSOCIATION) triples. **(b)** Global MRR: Model performance across all relations in the test set. Performance shown with original indications triples (blue) and degree preserving permutation for indication triples (red).

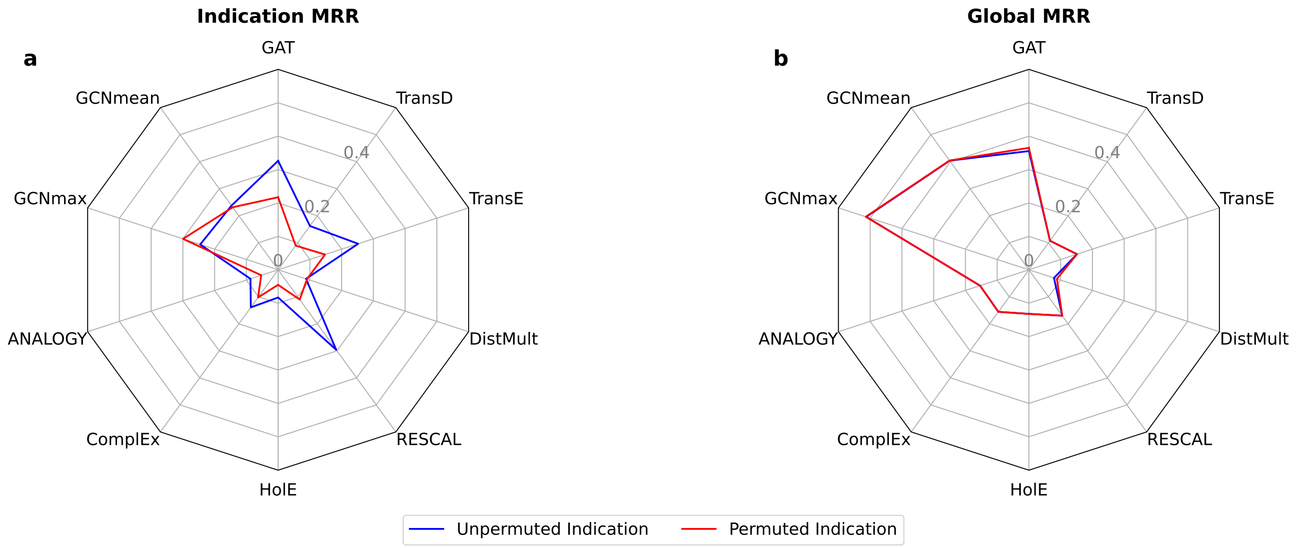


**Supplementary Figure S10.** Assessment of node degree exploitation as an illegitimate feature (DL2) in indication prediction on **HetionetKG**. **(a)** MRR for the Indication Relation: Model performance on indication (compound_treats_disease) triples. **(b)** Global MRR: Model performance across all relations in the test set. Performance shown with original indications triples (blue) and degree preserving permutation for indication triples (red).

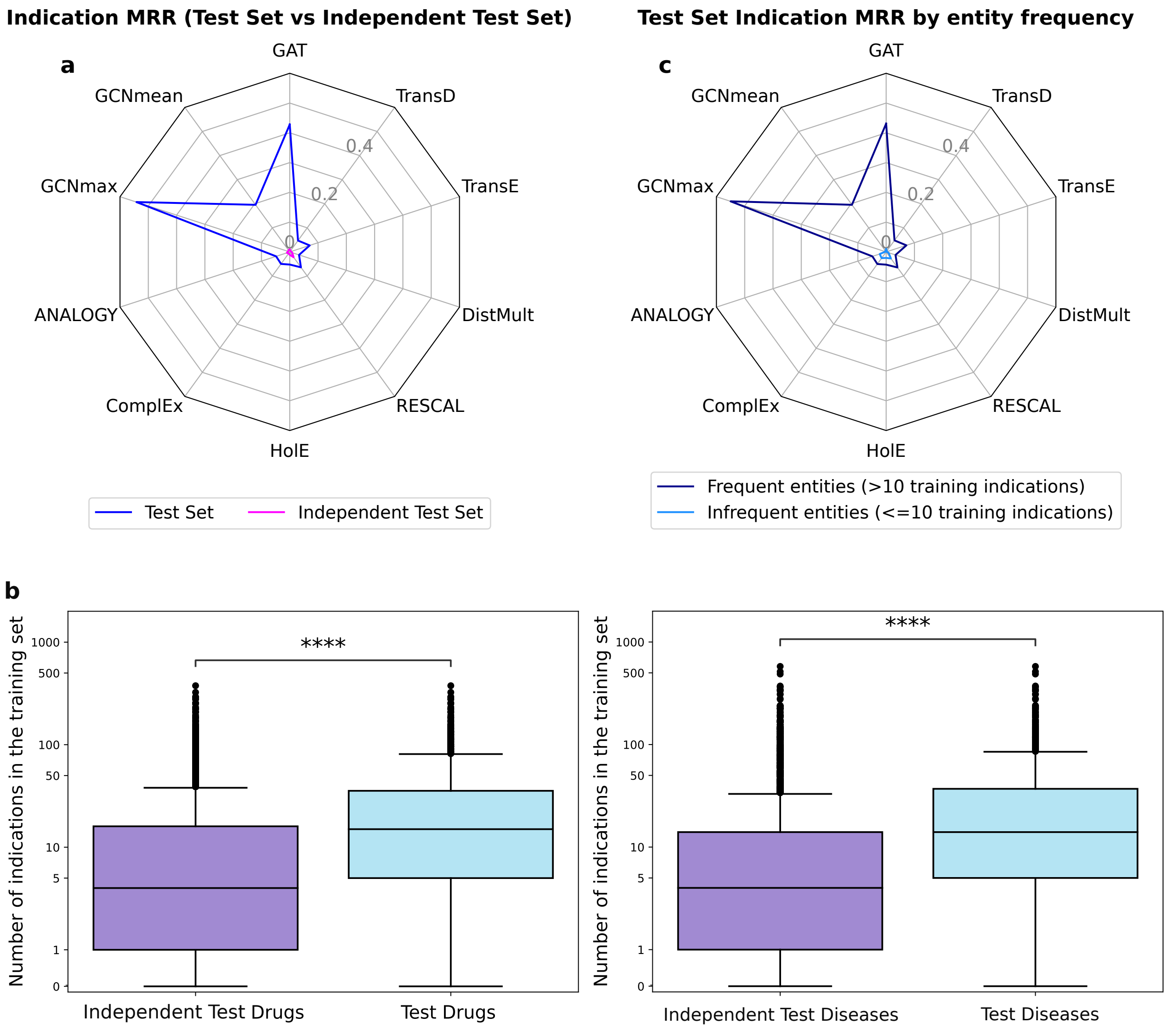


**Supplementary Figure S11.** Random Split Test Set and Independent Test Set Comparisons (DL3) on **BioKG**. **(a)** MRR comparison between test set of randomly sampled indication (DRUG_DISEASE_ASSOCIATION) triples (blue) and independent test set indication triples (pink). **(b)** Distribution of the number of training set indications (shown on a logarithmic scale) per drugs and per diseases across test and independent test sets. Statistical significance was assessed using the Mann-Whitney-Wilcoxon test (two-sided). Annotations indicate significance levels: ****: $p \leq 1.00 \times 10^{-4}$. **(c)** MRR comparison on test *indication* triples involving entities with frequent ($> 10$) or infrequent ($\leq 10$) occurrences in training *indication* triples (dark and light blue, respectively).

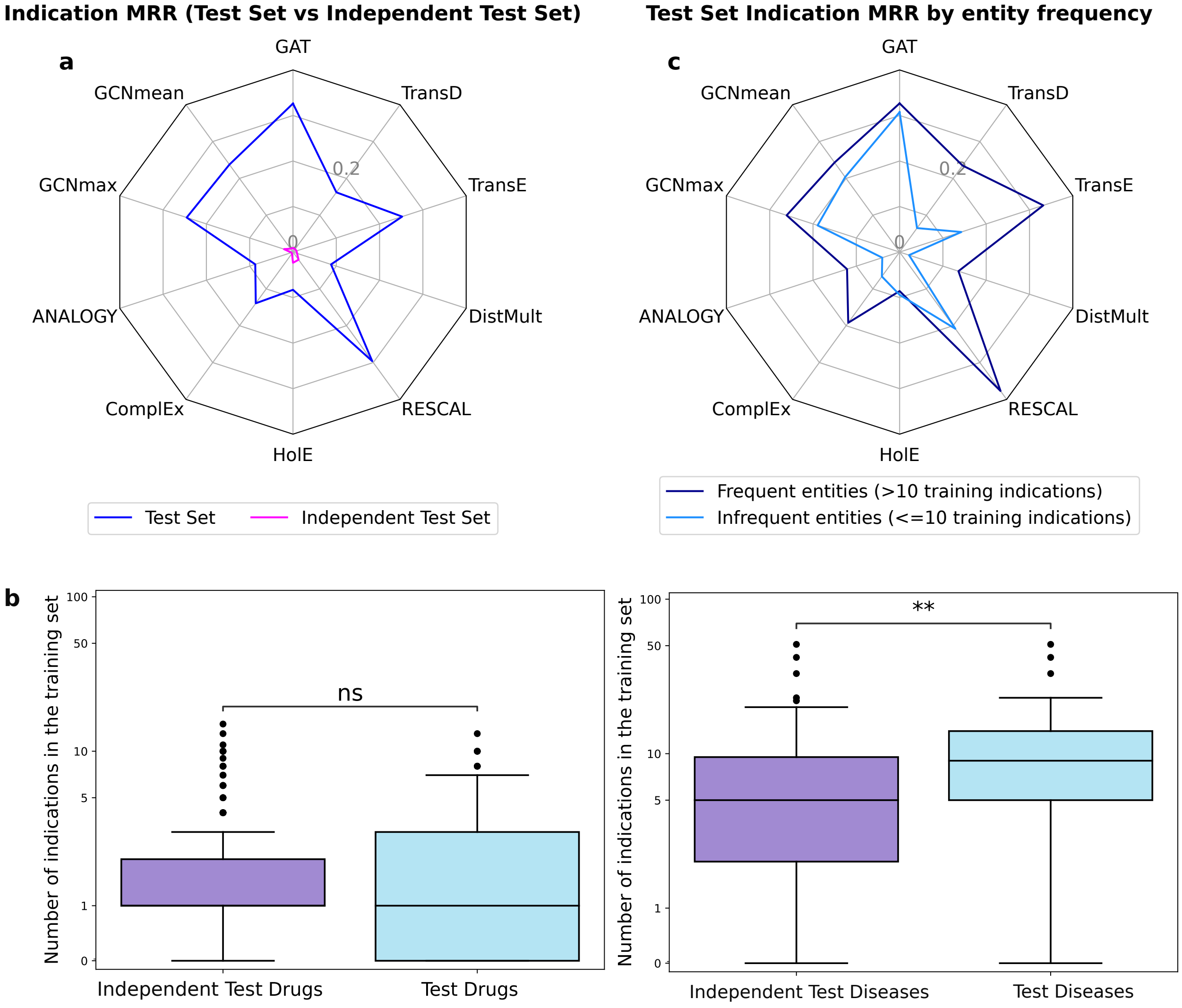


**Supplementary Figure S12.** Random Split Test Set and Independent Test Set Comparisons (DL3) on **HetionetKG**. **(a)** MRR comparison between test set of randomly sampled indication (compound_treats_disease) triples (blue) and independent test set indication triples (pink). **(b)** Distribution of the number of training set indications (shown on a logarithmic scale) per drugs and per diseases across test and independent test sets. Statistical significance was assessed using the Mann-Whitney-Wilcoxon test (two-sided). Annotations indicate significance levels: *ns*: $5.00 \times 10^{-2} < p \leq 1$; ****: $1.00 \times 10^{-3} < p \leq 1.00 \times 10^{-2}$. **(c)** MRR comparison on test *indication* triples involving entities with frequent ($> 10$) or infrequent ($\leq 10$) occurrences in training *indication* triples (dark and light blue, respectively).

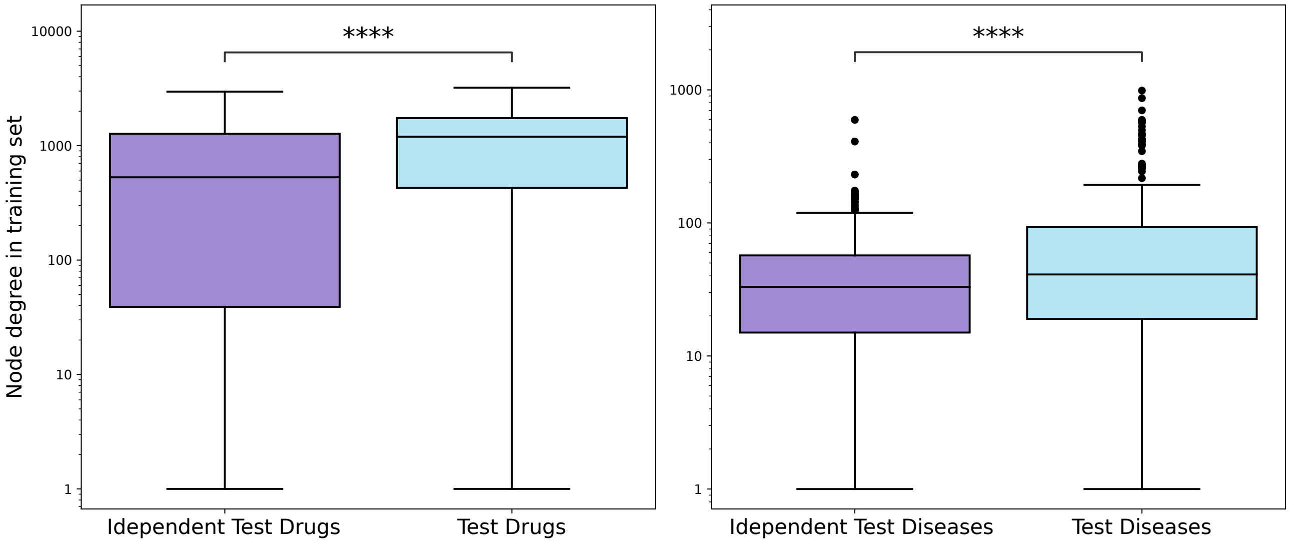


**Supplementary Figure S13.** Distribution, in **ShepherdKG**, of the total node degree in the training set (i.e., number of triples involving each entity across all relation types) for drugs and diseases from the test and independent test sets, shown on a logarithmic scale. Statistical significance was assessed using the Mann-Whitney-Wilcoxon test (two-sided). Annotations indicate significance levels: **** ($P \leq 1 \times 10^{-4}$).

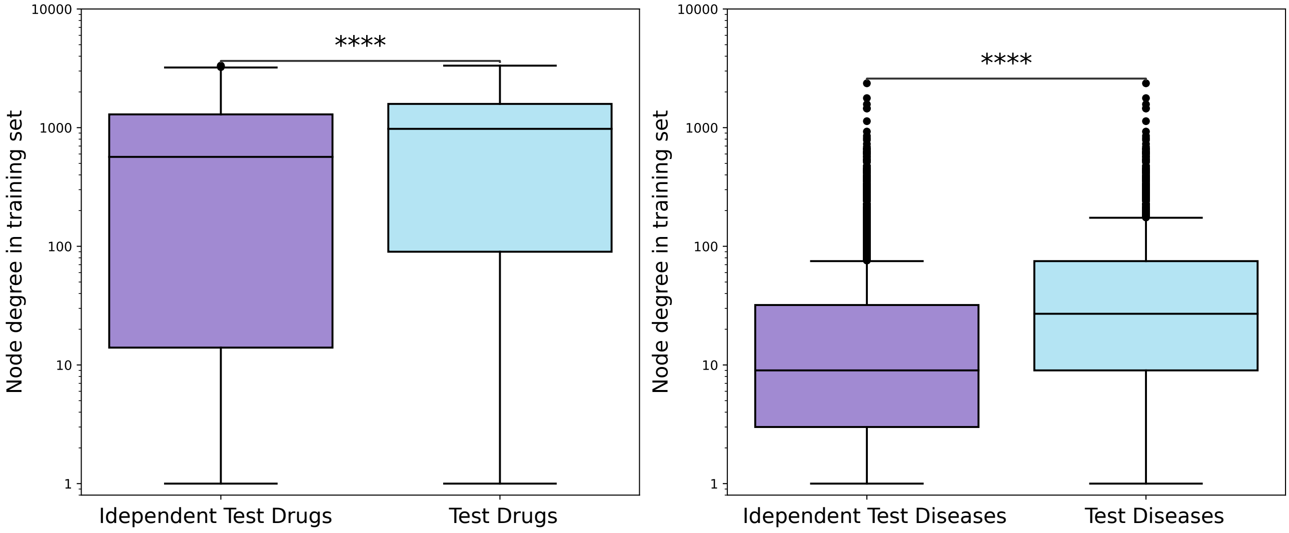


**Supplementary Figure S14.** Distribution, in **BioKG**, of the total node degree in the training set (i.e., number of triples involving each entity across all relation types) for drugs and diseases from the test and independent test sets, shown on a logarithmic scale. Statistical significance was assessed using the Mann-Whitney-Wilcoxon test (two-sided). Annotations indicate significance levels: **** ($P \leq 1 \times 10^{-4}$).

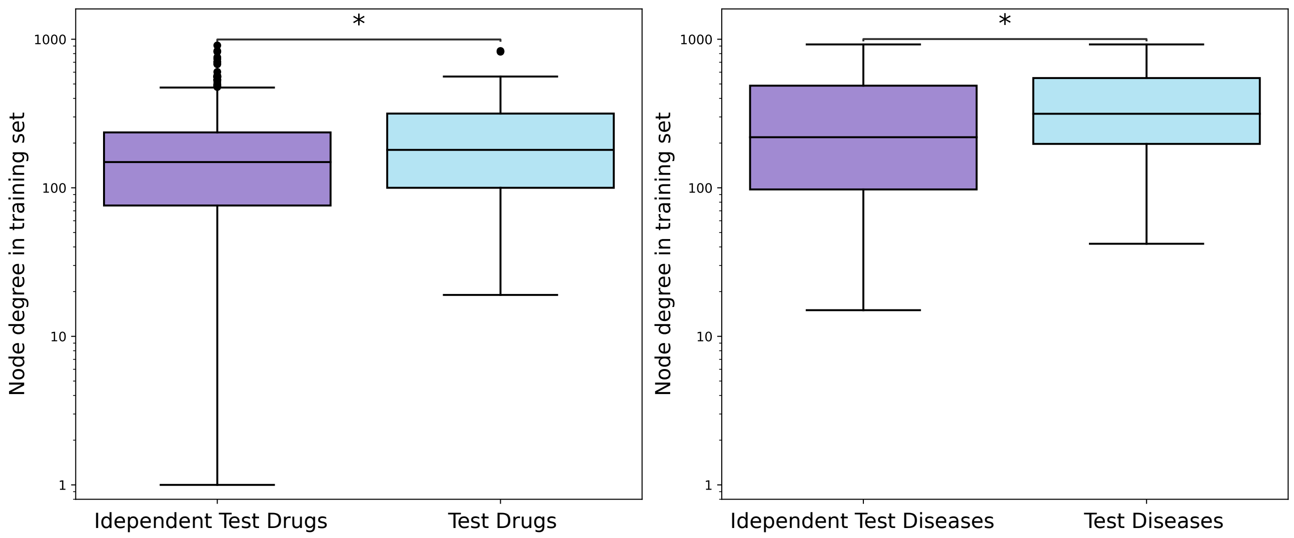


**Supplementary Figure S15.** Distribution, in **HetionetKG**, of the total node degree in the training set (i.e., number of triples involving each entity across all relation types) for drugs and diseases from the test and independent test sets, shown on a logarithmic scale. Statistical significance was assessed using the Mann-Whitney-Wilcoxon test (two-sided). Annotations indicate significance levels: *: $1.00 \times 10^{-2} < p \leq 5.00 \times 10^{-2}$.

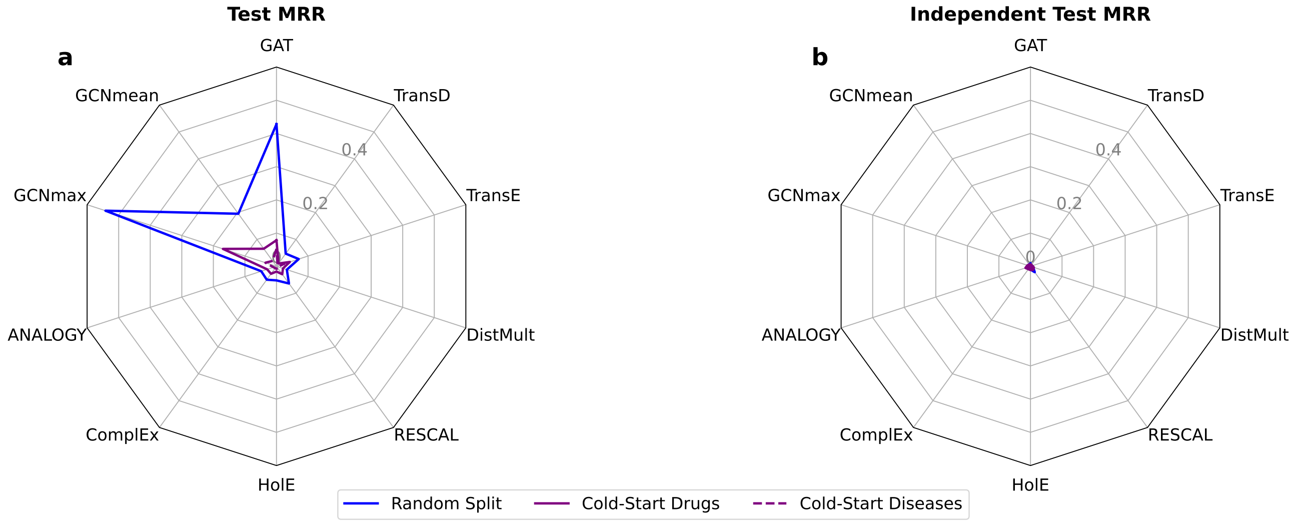


**Supplementary Figure S16.** Evaluating Test Set and Independent Test Set Performance under Random vs. Cold-Start Splits (DL3) on **BioKG**. **(a)** MRR comparison on the test set: randomly sampled indication triples (blue), drug-based cold-start indication triples (solid dark purple) and disease-based cold-start indication triples (dashed dark purple). **(b)** MRR comparison on the independent test set: randomly sampled indication triples (blue), drug-based cold-start indication triples (solid dark purple) and disease-based cold-start indication triples (dashed dark purple).

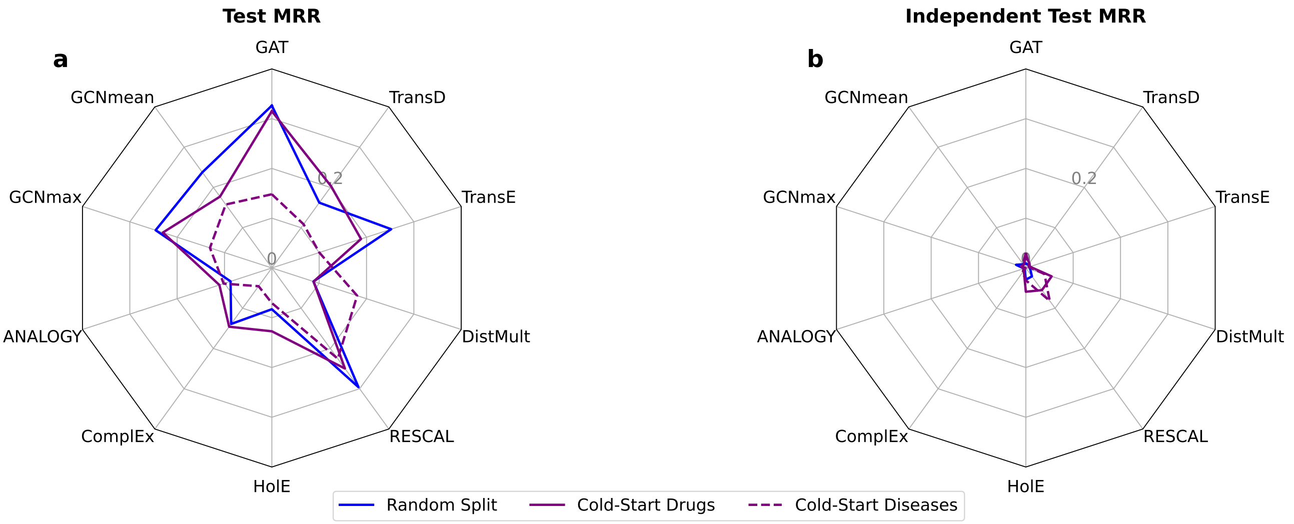


**Supplementary Figure S17.** Evaluating Test Set and Independent Test Set Performance under Random vs. Cold-Start Splits (DL3) on **HetionetKG**. **(a)** MRR comparison on the test set: randomly sampled indication triples (blue), drug-based cold-start indication triples (solid dark purple) and disease-based cold-start indication triples (dashed dark purple). **(b)** MRR comparison on the independent test set: randomly sampled indication triples (blue), drug-based cold-start indication triples (solid dark purple) and disease-based cold-start indication triples (dashed dark purple).

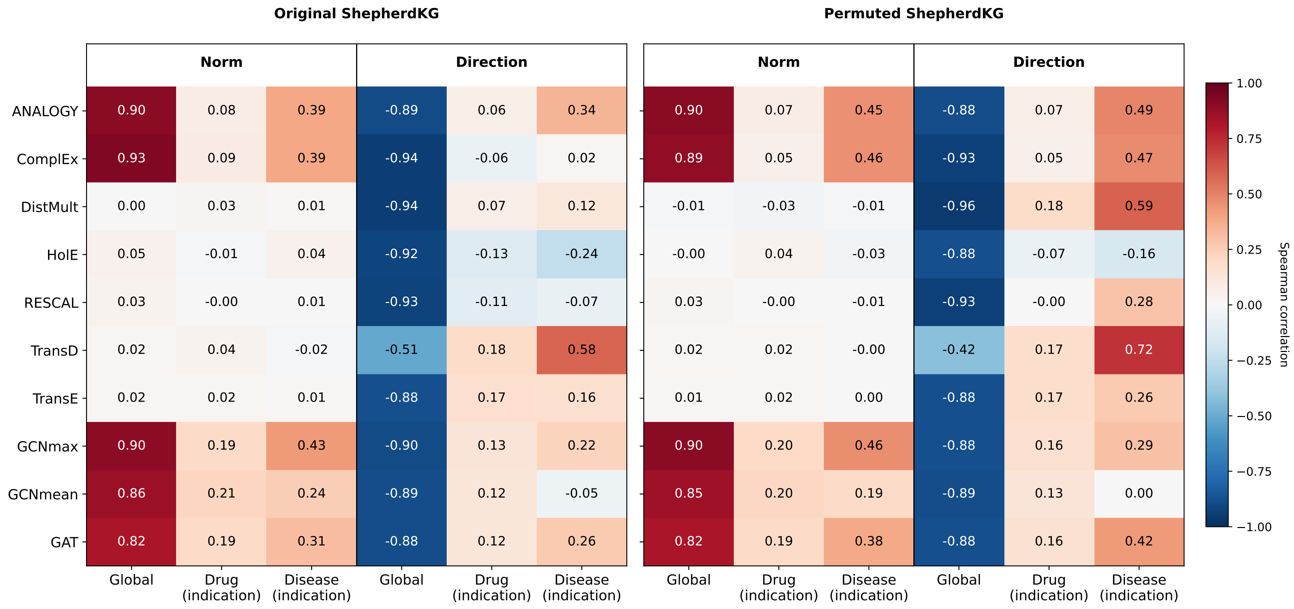


**Supplementary Figure S18.** Spearman correlations between embedding geometry and node degree on **ShepherdKG**. Each panel reports, for every model, the correlation between embedding geometry and node degree: the *Norm* block uses the $L_2$ norm of the entity embedding; the *Direction* block uses the cosine alignment of the unit embedding to the mean direction (centroid) of the relevant node set. Degree is defined at three levels: global (all relations, in + out, over all nodes) and, within the indication relation, the degree of drugs (heads) and diseases (tails), each computed over its own drug or disease node set. **Left:** original ShepherdKG; **Right:** degree-preserving DL2 permutation. Each correlation is averaged over two runs of the model (Main Text, Materials and Methods).

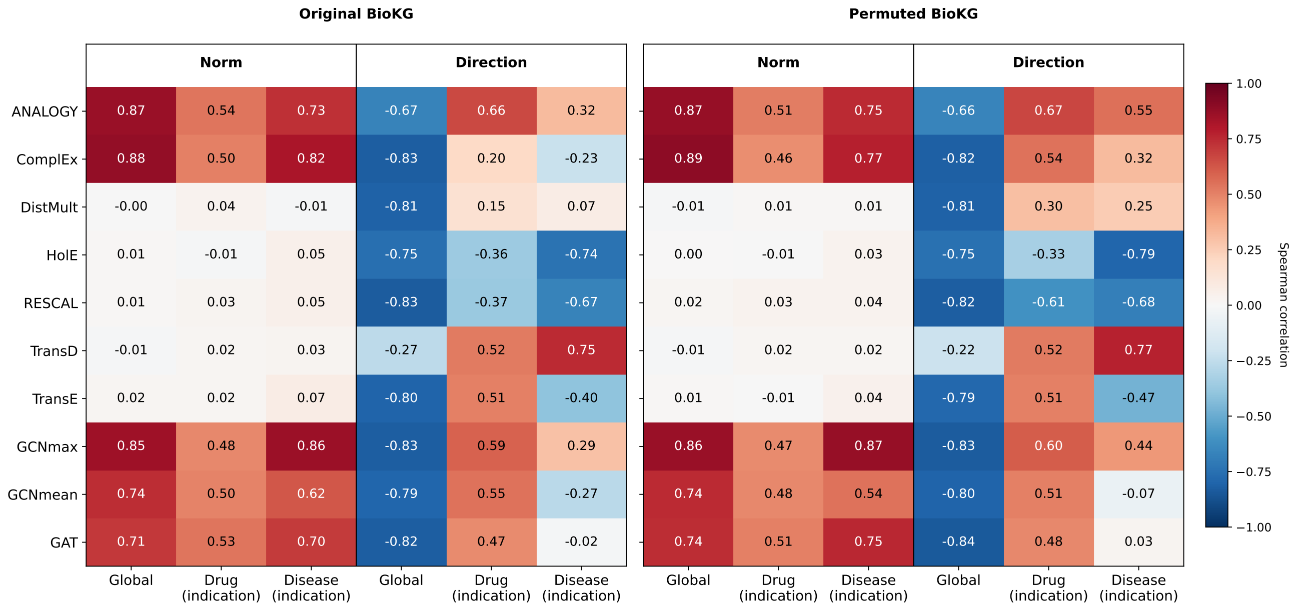


**Supplementary Figure S19.** Spearman correlations between embedding geometry and node degree on **BioKG**. Each panel reports, for every model, the correlation between embedding geometry and node degree: the *Norm* block uses the $L_2$ norm of the entity embedding; the *Direction* block uses the cosine alignment of the unit embedding to the mean direction (centroid) of the relevant node set. Degree is defined at three levels: global (all relations, in + out, over all nodes) and, within the indication relation (DRUG_DISEASE_ASSOCIATION), the degree of drugs (heads) and diseases (tails), each computed over its own drug or disease node set. **Left:** original ShepherdKG; **Right:** degree-preserving DL2 permutation. Each correlation is averaged over two runs of the model (Main Text, Materials and Methods).
